# Bacterial Vaginosis Toxins Impair Sperm Capacitation and Fertilization

**DOI:** 10.1101/2025.03.01.640991

**Authors:** Shweta Bhagwat, Leila Asadi, Ronald McCarthy, Juan Ferreira, Ping Li, Ethan Li, Sariela Spivak, Ariana Gaydon, Vaka Reddy, Christy Armstrong, Sydney R. Morrill, Hillary Zhou, Amanda L. Lewis, Warren G. Lewis, Celia M. Santi

**Affiliations:** Department of Obstetrics and Gynecology, Washington University School of Medicine, Saint Louis, MO, USA; Department of Obstetrics, Gynecology, and Reproductive Sciences, Glycobiology Research and Training Center, University of California San Diego, La Jolla, California, USA

**Keywords:** Sperm capacitation, hyperactivation, acrosomal exocytosis, infertility, bacterial vaginosis, lipopolysaccharide, vaginolysin, CatSper, toll-like receptor 4

## Abstract

**Study question:** What effect do toxins produced by bacterial vaginosis (BV) bacteria have on sperm function?

**Summary answer:** Bacterial vaginosis toxins dysregulate sperm capacitation and intracellular calcium homeostasis and impair the ability of sperm to fertilize oocytes.

**What is known already:** In bacterial vaginosis, which is linked to infertility, overgrowth of *Prevotella* and *Gardnerella* in the vagina is accompanied by elevated concentrations of the toxins lipopolysaccharide (LPS) and vaginolysin (VLY).

**Study design, size, duration:** This was a laboratory study in which human semen samples were collected from consenting healthy donors with normal semen parameters. Mouse sperm samples were obtained from the caudal epididymis.

**Participants/materials, setting, methods:** Motile mouse and human sperm were isolated via swim-up and treated under non-capacitating or capacitating conditions. LPS from *Escherichia coli* was commercially available. VLY was produced by cloning the *Gardnerella* VLY protein in the ClearColi expression system. Mouse sperm were pre-incubated in *in vitro* fertilization medium with LPS or VLY and then co-cultured with ovulated cumulus-oocyte complexes. The effects of LPS and VLY on sperm motility and hyperactivation were assessed with computer-assisted sperm analysis. Effects on viability were assessed by Hoechst staining. Acrosomal exocytosis was assessed in sperm from transgenic Acr-eGFP mice and in human sperm stained with *Pisum sativum* agglutinin FITC. Intracellular calcium dynamics were assessed by staining sperm with the calcium-sensitive dye Fluo-4 AM and fluorescent imaging several sperm at the single-cell level. The effects of LPS on sperm from CatSper knock-out mice were assessed. Additionally, sperm were treated with a toll-like receptor 4 antagonist and further exposed to LPS.

**Main results and the role of chance:** Exposure of mouse sperm to LPS or VLY significantly decreased *in vitro* fertilization (*P* < 0.05). Under capacitating conditions, both toxins initially increased mouse and human sperm hyperactivation, then significantly decreased sperm motility (*P* < 0.05), hyperactivation (*P* < 0.05), and acrosomal exocytosis (*P* < 0.01). These changes were accompanied by a rapid and irreversible increase in intracellular calcium concentration. Effects of LPS, but not VLY, were prevented by polymyxin-B, which aggregates LPS. The LPS-induced intracellular calcium increase required external calcium but not the calcium channel CatSper and was inhibited by the Toll-like receptor 4 antagonist.

**Limitations, reasons for caution:** First, the commercially available LPS we used was isolated from *Escherichia coli*, rather than from the BV-associated bacteria *Prevotella bivia*. Second, we did not quantify the absolute sperm intracellular calcium concentration before or after LPS or VLY treatment. Third, all of our experiments were *in vitro*.

**Wider implications of the findings:** These studies suggest that BV-associated toxins contribute to infertility by, in part, impairing sperm capacitation and reducing their fertilizing ability.

**Study funding/competing interest(s):** This work was supported by the National Institutes of Health (grant #R01 HD069631). The authors declare that they have no conflict of interest.

## Introduction

Bacterial vaginosis (BV) is characterized by a disruption of the vaginal microbiota, with a decrease in protective *Lactobacillus* species and overgrowth of anaerobic bacteria, including *Gardnerella* and *Prevotella* (Ravel *et al*., 2011). BV affects approximately 29% of reproductive-aged women in the U.S. (Allsworth and Peipert, 2007), and is associated with various adverse health outcomes, such as sexually transmitted infections, pelvic inflammatory disease (Ravel *et al*., 2021; Turpin *et al*., 2021), and preterm birth (Salminen *et al*., 2008). Moreover, BV has been linked to female infertility through several mechanisms (Ravel *et al*., 2021; Cocomazzi *et al*., 2023). First, elevated concentrations of pro-inflammatory cytokines and chronic inflammation in the reproductive tract lead to tissue damage and impaired implantation (Murphy and Mitchell, 2016; Ravel *et al*., 2021). Second, BV-associated bacteria and their metabolites induce endometrial dysfunction (Łaniewski and Herbst-Kralovetz, 2021), interfere with embryo implantation, and cause early pregnancy loss (Eckert *et al*., 2003; Oostrum *et al*., 2013; Haahr *et al*., 2018; Hong *et al*., 2020). Third, imbalances in vaginal pH and microbiota (Criteria and Associations, 1983; Mania-pramanik *et al*., 2008), along with the toxins produced by *Gardnerella* and *Prevotella*, adversely affect sperm quality and motility (Li *et al*., 2016; O’Doherty *et al*., 2016).

*Prevotella* is an opportunistic pathogen in the female genital tract (George *et al*., 2024). Like other *Proteobacteria,* it produces lipopolysaccharide (LPS) (Aroutcheva *et al*., 2008), a heat-stable toxin (also known as endotoxin). LPS reaches 2-3 orders of magnitude higher concentrations in the vagina of those with BV than in those without BV. These higher LPS concentrations strongly correlate with the abundance of *Prevotella* in BV-positive individuals (Aroutcheva *et al*., 2008). LPS-mediated inflammation has been linked with female infertility, affecting ovarian and pituitary function (Magata *et al*., 2023). Additionally, in several animal models, LPS challenge can trigger preterm birth and injure the fetal brain (Wang *et al*., 2006; Firmal *et al*., 2020). Unlike *Prevotella*, *Gardnerella* does not make LPS but instead produces the toxin vaginolysin (VLY). VLY is a cholesterol-dependent cytolysin that forms pores in human cell membranes and is active against erythrocytes, as well as vaginal and cervical cells *in vitro* (Yoo and Lee, 2016; Ragaliauskas *et al*., 2019; Campisciano *et al*., 2021).

We hypothesize that LPS and VLY contribute to female infertility by disrupting sperm capacitation, which occurs in the female genital tract and is necessary for a sperm to be able to fertilize an egg. One component of capacitation is hyperactivation, which causes flagellar beating to increase in amplitude, decrease in frequency, and become asymmetrical (Suarez, 2008). Capacitation also involves acrosomal exocytosis (Molina *et al*., 2018) in which the outer acrosomal membrane at the sperm head fuses with the sperm plasma membrane to release the acrosomal content in response to a stimulus (Sosa *et al*., 2014). Hyperactivation and acrosomal exocytosis are necessary for sperm to penetrate the cumulus and zona pellucida layers surrounding the oocyte and fuse with the oocyte.

Here, we show that both LPS and VLY caused mouse and human sperm to hyperactivate early during capacitation. Prolonged exposure to LPS and VLY decreased sperm motility, hyperactivation, and fertilization capability. Furthermore, sperm acrosomal exocytosis was inhibited by the two toxins. In defining the underlying mechanisms, we found that LPS and VLY caused rapid and irreversible increases in intracellular calcium (Ca^2+^) concentration in sperm. Although hyperactivation and acrosomal exocytosis require an increase in intracellular Ca^2+^, excessive intracellular Ca^2+^ concentration impairs mitochondrial function, leading to a loss of motility, activation of apoptotic cascades, and premature loss of the acrosome, in mammalian sperm (Liu and Baker, 1990). Finally, we show that the LPS-induced Ca^2+^ entry involved Toll-like receptor 4 (TLR4). VLY-induced Ca^2+^ entry is likely due to the membrane pores formed by this toxin. Together, our data show that toxins present in BV dysregulate capacitation and intracellular Ca^2+^ homeostasis, impairing the ability of sperm to fertilize oocytes. This mechanism may contribute to infertility in patients with BV.

## Materials and methods

### Reagents

All reagents used to prepare Human Tubal Fluid (HTF) and Toyoda–Yokoyama–Hosi (TYH) media, as well as the Ca^2+^ ionophore A23187, polymyxin B (PMB), TAK-242 (TLR4 receptor inhibitor), progesterone, Methyl-ß-cyclodextrin, and immunoglobulin-free bovine serum albumin (BSA) were from Millipore Sigma (St. Louis, MO, USA). Dulbecco’s Phosphate Buffered Saline (DPBS) tissue culture grade was from GIBCO Life Technologies (Gaithersburg, MD, USA); Hoechst 33342 was from Cayman Chemicals (Ann Arbor, MI, USA); Fluo-4 AM was from Invitrogen^TM^ Thermo Fisher Scientific (Waltham, MA, USA); paraformaldehyde was from Electron Microscopy Sciences (Hatfield, PA, USA); and LPS from *E. coli* strain O112:H10 was from Associates of Cape Cod Inc. (E. Falmouth, MA, USA).

### Animals and ethics statement

All mouse procedures were performed according to the National Institutes of Health Guiding Principles for the care and use of laboratory animals. These procedures were reviewed and approved by the Institutional Animal Care and Use Committee of Washington University in St. Louis (St. Louis, MO, USA) (protocol # 20-0126).

C57BL/6 and CatSper1 knock-out male mice (CatSper KO) (60–90 days old) were purchased from Jackson Labs and kept at a constant temperature of 22 ± 2 °C under a 12/12 h dark/light cycle with free access to food and water. Oocytes for *in vitro* fertilization experiments were obtained from 4–8-week-old C57BL/6 female mice.

### Mouse sperm collection and capacitation

Sexually mature male mice (60–90 days old) were euthanized via cervical dislocation, and sperm were isolated from the cauda epididymis. Sperm were allowed to swim-up in non-capacitating (NC) TYH media buffered with HEPES (in mM: 119.3 NaCl, 4.7 KCl, 1.71 CaCl_2_.2H_2_O, 1.2 KH_2_PO_4_, 1.2 MgSO_4_.7H_2_O, 25.1 NaHCO_3_, 5.56 glucose, 0.51 sodium pyruvate, 10 HEPES) at pH 7.4 for 15–20 min at 37 °C, unless specified. The motile fraction of the sample was removed from the tube. To induce capacitation, sperm were incubated in capacitating (CAP) TYH medium supplemented with 5 mg/mL BSA and 15 mM NaHCO_3_ at 37 °C for 60-90 min. NC sperm were incubated in TYH without BSA and NaHCO_3_ unless specified otherwise.

### Human participants and sperm preparation

Human semen samples were obtained by masturbation after 3 to 5 days of abstinence. Samples were left over from the Washington University Fertility and Reproductive Medicine Center and were deidentified. The clinic confirmed that all samples met normal semen parameters according to the World Health Organization (≥15 million sperm per mL, ≥40% total motility, ≥32% progressive motility). Samples were allowed to liquefy for 1 h at room temperature, then sperm were purified by swim-up as previously described (Molina *et al*., 2020). Briefly, within 2 h of production, sperm were allowed to swim up in non-commercial HTF (in mM: 98 NaCl, 4.7 KCl, 0.4 KH_2_PO_4_, 2 CaCl_2_, 0.2 MgSO_4_, 20 HEPES, 3 glucose, 21 lactic acid, 0.3 sodium pyruvate; pH adjusted to 7.4 with NaOH) for 1 h at 37 °C without CO_2_.

### Production of *Gardnerella* vaginolysin (VLY) protein

A truncated mutant *vly* gene from *Gardnerella* ATCC14019 (Randis *et al*., 2009) was cloned into a pET28a vector, which was transformed into ClearColi BL21(DE3) cells (Biosearch Technologies, Teddington, UK) with detoxified LPS. Protein was expressed and purified as previously described (Morrill *et al*., 2023). Cytolytic activity on human red blood cells was verified by serial dilution and assay of hemoglobin release measured at 545 nm. In parallel, an empty vector was used to prepare a mock expression and purification that lacked VLY but contained all other reagents. VLY and mock preparations were stored in 20% glycerol at -20 °C.

### LPS precautions and testing

LPS from Associates of Cape Cod, Inc. (E Falmouth, MA) was dissolved according to the manufacturer’s instructions. Dilutions were performed in a manner that avoided loss of LPS, avoiding multiple dilutions in plastic that would allow adsorption. Because LPS can contaminate reagents, especially recombinant proteins, pyrogen-free reagents and labware were used when available. Clean glassware was dry baked at 200 °C for 2 h to destroy LPS. The LPS content of reagents was quantified by using a chromogenic Limulus Amoebocyte Lysate assay from Associates of Cape Cod. Protein preparations and buffers (TYH and HTF) contained <1 EU/mL (approximately <0.1 ng/mL) of LPS at the concentrations used in sperm preparation and assays.

### *In vitro* fertilization (IVF) protocol

Sperm from C57BL/6 male mice (4–8 weeks old) were prepared as described above and capacitated for 3 h in TYH-CO_2_ buffered media supplemented either with 0.75 mM Methyl-ß-cyclodextrin (Nakagata, 2010) or 5 mg/mL BSA at 37 °C and 5 % CO_2_ to induce capacitation, with or without LPS or VLY. C57BL/6 female mice (4–8 weeks old) were superovulated by intraperitoneally injecting 5-7.5 IU pregnant mare serum gonadotropin (ProSpec Cat# HOR-272). 48 h later, these mice were injected with 5-7.5 IU human chorionic gonadotropin (Millipore/Sigma CG5-1VL). 12-15 h later (at the 3-h sperm capacitation timepoint), female mice were euthanized, and oocyte-cumulus complexes were isolated in 200 µL of high Ca^2+^ HTF (Nakagata, 2010). Cumulus-oocyte complexes were washed four times with 100 µl HTF, then overlayed with Ovoil. The media in the dish was allowed to equilibrate for at least 1 h in an incubator at 37 °C and 5% CO_2_, before the addition of oocytes. Fertilization wells containing 20–30 eggs were mixed with CAP sperm (final concentration of 2 × 10^6^ sperm/mL) and incubated for 4-6 h. Eggs were then washed three times and placed in a drop of HTF medium with a disposable soda-lime glass 50 µL microcapillary pipette (Kimble Cat# 71900-50) attached to an aspirator tube assembly (Millipore/Sigma A5177-5EA). Oocytes were counted, and the dish was incubated overnight at 37 °C and 5% CO_2_. The percentage of fertilized eggs was calculated by dividing the number of two-cell embryos by the total number of oocytes.

### Computer-Assisted Sperm Analysis (CASA)

Mouse and human sperm were suspended in their respective CAP media and exposed to 1 µg/mL LPS or VLY (mouse sperm) and 0.1 µg/mL LPS or VLY (human sperm) from the start of capacitation. Sperm motility and hyperactivity were measured at time 0 and at 0.5-1 h intervals. Sperm samples (3 μl) from each condition were placed into 20-micron Leja standard count four-chamber slides, pre-warmed at 37 °C, and a minimum of 200 cells were counted. CASA was performed with a Hamilton–Thorne digital image analyzer (HTR-CEROS II v.1.7; Hamilton–Thorne Research, Beverly, MA, USA). CASA settings were as previously described (Molina *et al*., 2020). For mouse sperm, hyperactivated motility was defined as curvilinear velocity (VCL) > 150 μm/s, lateral head displacement (ALH) > 7.0 μm, and linearity coefficient (LIN) of 32% at 60 Hz.(Mortimer *et al*., 1998) For human sperm, hyperactivated motility was defined as VCL ≥ 150 μm/s, ALH ≥ 7 μm, LIN ≤ 50% at 60 Hz. Percent hyperactivation was calculated by dividing the number of hyperactivated sperm by the number of motile sperm. Total and progressive motility values were obtained as percentages from the CASA software.

### Sperm viability assessment

Hoechst 33342 (900 nM) was added to sperm. After 2 to 5 min, fluorescence intensity was measured with an Aurora 4L 16V-14B-10YG-8R Flow Cytometer and a V3 or 458 nm filter (Lyon *et al*., 2023). The excitation wavelength of Hoechst 33342 is 350 nm, and its emission wavelength is 461 nm. At least 10,000 sperm were recorded for each sample. The fluorescence cutoff was selected by measuring the fluorescence of sperm killed by treatment with 3 to 6% sodium hypochlorite for 2 to 5 min.

### Sperm intracellular calcium concentration measurement

After sperm swim-up in NC or CAP conditions for 15–20 min, motile sperm were collected and incubated with 10 μM Fluo-4 AM and 0.05% Pluronic F-127 at 37 °C for 60 min. Sperm were centrifuged at 300-400 g for 5–10 min, resuspended in the corresponding media, and allowed to attach to Poly-l-lysine (0.1%) coated coverslips for 5 min. A local perfusion device with an estimated exchange time of 10 s was used to apply test solutions.

Sperm [Ca^2+^]_I_ was recorded at room temperature with a Leica AF 6000LX system connected to a Leica DMi8000 inverted microscope, equipped with a 63X objective (HC PL FluoTar L 63X/0.70 Dry) and an Andor-Zyla-VCS04494 camera. A halogen lamp was used with a 488 ± 20 nm excitation filter and a 530 ± 20 nm emission filter. Data were collected with Leica LasX 2.0.014332 software. Acquisition parameters used were: 20 ms exposure time, 4x4 binning, 1024 x 1024 pixels resolution (Ferreira *et al*., 2021a). For each condition, an initial baseline reading was followed by the addition of LPS or VLY and a final perfusion with 5 μM Ionomycin (Iono) at the end of every experiment. Whole images were collected every 10 s. LAS X, ImageJ, Clampfit 10 (Molecular Devices), and GraphPad were used for data analysis. A region of interest was selected in the sperm head (Chávez *et al*., 2014). To compare fluorescence across different samples and experimental conditions, Fluo-4 fluorescence (F) in response to LPS or VLY for each sperm was normalized to its respective F_Iono_ (F/F_iono_), after background subtraction. Cells with [Ca^2+^]_I_ increases of >10% of that obtained with Iono were counted as responsive. Percentages of cells responding to LPS or VLY were calculated by dividing the number of cells responding to the respective toxin by the total number of cells that responded to Iono. Clampfit 10 software was used to measure amplitude and Tau (time taken to exhibit the response) in responding sperm (Ferreira *et al*., 2021a).

### Acrosomal exocytosis

For mouse sperm, acrosomal exocytosis was evaluated in both NC and CAP sperm from transgenic mice that express green fluorescent protein in the acrosome (Hasuwa *et al*., 2010). Acrosomal exocytosis was induced by incubating sperm with 15 µM Ca^2+^ ionophore A23187 or 50 mM KCl (Millipore Sigma, St. Louis) for 30 min before the end of capacitation. LPS or VLY (1 µg/mL) were added to CAP and NC samples either at the start of capacitation or with the inducers A23187 and KCl. Sperm from all conditions were fixed in 4% paraformaldehyde and centrifuged at 300-400 g for 5–10 min. Supernatants were removed, the sperm were washed with phosphate buffered saline (PBS), and 15 μL of each sample was smeared onto a glass slide. Slides were air-dried and coverslips were mounted with 1:1 glycerol: pyrogen-free DPBS at room temperature. Sperm were examined at 40X on an EVOS FL Cell Imaging System epifluorescence microscope (Thermo Fischer Scientific). Sperm were counted as acrosome-reacted if the acrosomal region lacked GFP fluorescence and non-reacted if they retained GFP fluorescence. A minimum of 200 sperm were evaluated in each experiment. Percent acrosomal exocytosis was calculated by dividing the number of reacted sperm by the number of total sperm per sample.

Human sperm acrosomal exocytosis was measured as previously described (Lyon *et al*., 2023). Briefly, swim-up sperm were incubated in both NC and CAP media (HTF supplemented with 25 mM HCO_3_ and 5 mg/mL BSA) for 3 h. Acrosomal exocytosis was induced by adding 10 µM Ca^2+^ ionophore A23187 or 10 µM progesterone for 30 min. LPS or VLY (0.1 µg/mL) were added immediately after swim-up or during acrosomal exocytosis induction. After induction, sperm were fixed with 3% paraformaldehyde in DPBS at 4 °C, smeared on slides, and stained with FITC-labelled *Pisum sativum* agglutinin (Millipore Sigma). Fluorescence was imaged as described for mouse sperm acrosomal exocytosis. Sperm with bright, uniformly stained acrosomes were counted as intact, and sperm with equatorial or no staining in the acrosomal region were counted as reacted. The minimum number of sperm and method for calculating percent acrosomal exocytosis were as described for mouse sperm.

### Statistical analyses

GraphPad Prism version 10.1.0 for Windows (GraphPad Software, La Jolla, CA) was used for all statistical analyses. An unpaired two-tailed Student’s t-test (alpha < 0.05) was used to compare sperm samples from different individuals, whereas a paired two-tailed t-test was used to compare control and treatment in the same sample, unless otherwise indicated. For time series data analysis, where time and the treatment groups were the two variables, two-way ANOVA with Bonferroni’s multiple comparison test was performed. Data are expressed as the mean ± SD. *P* < 0.05 was considered statistically significant.

## Results

### LPS and VLY interfere with mouse sperm fertility and capacitation

To determine whether LPS or VLY impair the ability of sperm to fertilize oocytes, we capacitated mouse epididymal sperm in the presence and absence of 1 µg/mL LPS or 1 µg/mL VLY for 3 h. We then performed mouse *in vitro* fertilization. When sperm were capacitated in the presence of LPS, the fertilization rate was 34.93 ± 8.57%, significantly lower than the rate of 60.57 ± 4.28% in control conditions. Similarly, fertilization rates decreased from 76 ± 16% to 44 ± 16% in the absence and presence of VLY, respectively **(Fig. 1A, B)**. The addition of LPS or VLY to the fertilization drop after capacitation had no effect on fertilization rates (**Fig. S1** and **Fig. S2**).

**Figure 1:**
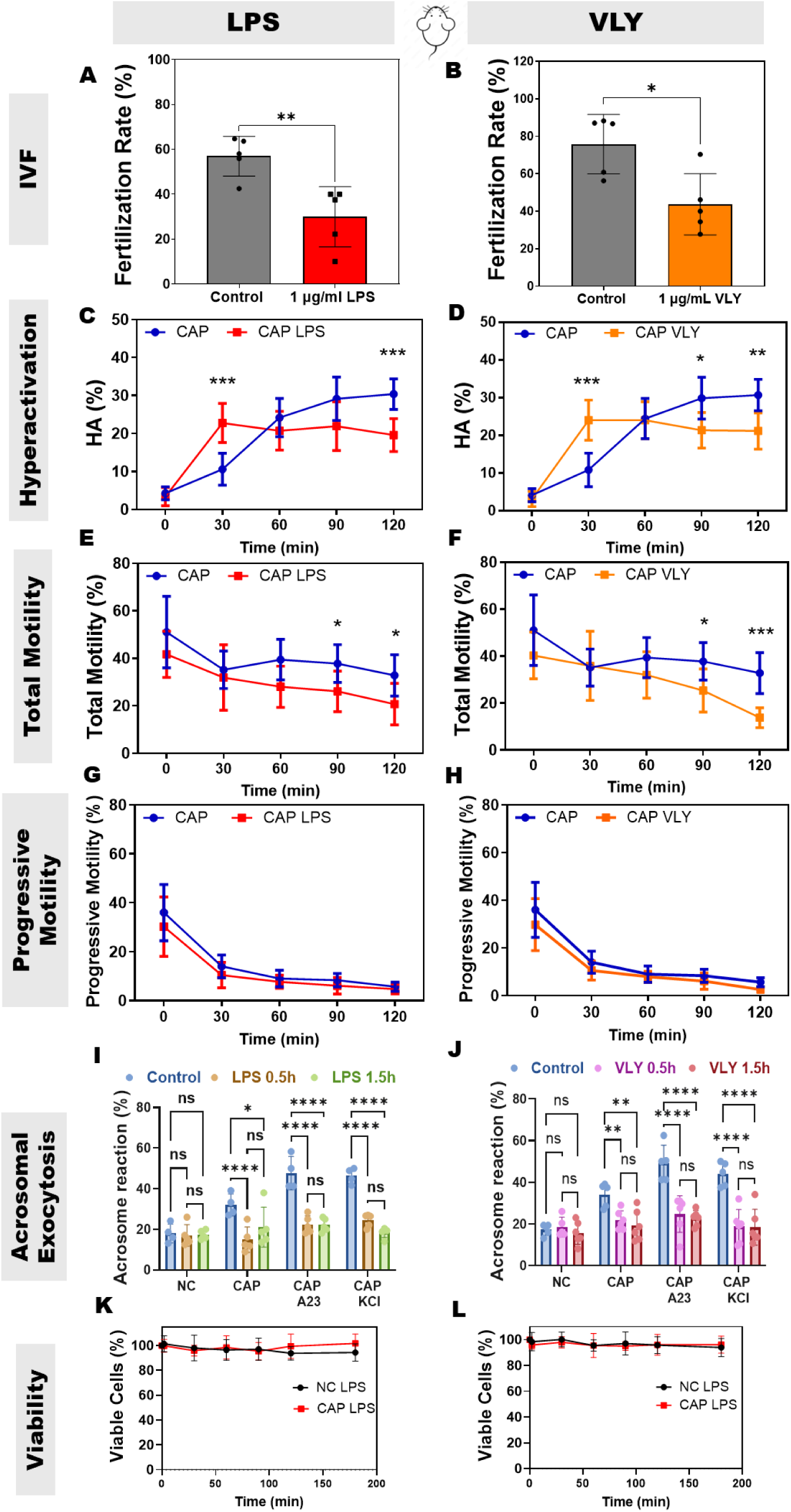
LPS and VLY impair mouse sperm fertility, hyperactivation, total motility, and acrosomal exocytosis in capacitating conditions. Percentage of oocytes that reached the two-cell stage within 24 h after *in vitro* fertilization (IVF) using sperm (n=5 biological replicates) previously treated for 3 h with 1 µg/mL **(A)** LPS or **(B)** VLY were calculated. CASA measurements were obtained for **(C, D)** hyperactivated motility, (**E, F**) total motility, and (**G, H**) progressive motility of mouse sperm incubated under capacitating (CAP) conditions in the presence and absence of 1 µg/mL (**C, E, G**) LPS and **(D, F, H)** VLY (n=9 biological replicates). Here 0 min is the time-point of BSA + sodium bicarbonate addition to initiate *in vitro* capacitation. Acrosomal exocytosis (AE) was quantified in non-CAP (NC), and CAP sperm (200 each, from 5 biological replicates) incubated with **(I)** LPS and **(J)** VLY, for 0.5 and 1.5 h. Similarly, AE induced by 15 µM A23187 or 50 mM KCl, was also quantified in CAP sperm. Sperm viability for NC and CAP sperm (n≥3 biological replicates) was analyzed in the presence of 1 µg/ml **(K)** LPS or **(L)** VLY (10,000 sperm each). Data are presented as mean and standard deviation. * P<0.05, ** P<0.01, *** P<0.001, **** P<0.001. For **A-B**, a paired t-test was used. For **C-L**, two-way ANOVA with Bonferroni’s multiple comparison test was used.

Next, we wanted to determine which outcomes of capacitation – hyperactivation, acrosomal exocytosis, or both – were affected by these toxins. Mouse sperm were incubated in capacitating (CAP) media with or without 1 µg/mL LPS or VLY, and motility was measured every 30 min for 120 min. At 30 min, the percentages of hyperactivated sperm were significantly higher in the presence of LPS or VLY than in the controls **(Fig. 1C, D)**. However, at 90 and 120 min, sperm hyperactivation and total motility were significantly lower in the presence of LPS or VLY than in controls (**Fig. 1C–F**). No effect was observed on progressive motility in CAP (**Fig. 1G, H**) or in non-CAP conditions (**Fig. S3**). These results suggest that LPS and VLY impair sperm hyperactivation, which is required for fertilization.

To measure the impact of LPS and VLY on acrosomal exocytosis, mouse sperm were exposed to 1 µg/mL LPS or VLY in CAP conditions, and acrosomal exocytosis was induced by adding 15 µM Ca^2+^ ionophore A23187 (A23) or 50 mM potassium chloride (KCl).(De La Vega-Beltran *et al*., 2012) As reported earlier (De La Vega-Beltran *et al*., 2012), about 40–60% of mouse sperm underwent induced acrosomal exocytosis in control CAP conditions. The presence of LPS or VLY significantly inhibited A23- and KCl-induced sperm acrosomal exocytosis but had no significant effect on acrosomal exocytosis in non-CAP conditions **(Fig. 1I, J)**, which is also defined as spontaneous acrosome reaction (Liu and Baker, 1990).

Next, we assessed whether these effects were specific to hyperactivation and acrosomal exocytosis or caused by a decrease in sperm viability. We found that mouse sperm viability did not significantly differ in the absence or presence of either toxin over 180 min (**Fig. 1K, L**). To test whether the effects were specifically due to LPS, we capacitated sperm in the presence of LPS plus polymyxin B (PMB), which forms aggregates with LPS and hampers its activity. In this condition, LPS had no effect on mouse sperm hyperactivation (**Fig. 2A**). PMB alone also had no effect on hyperactivation (**Fig. 2B**). As a control for VLY specificity, we treated sperm with protein generated in parallel using an empty vector control. This treatment did not impair hyperactivation (**Fig. 2C**). Additionally, the changes in hyperactivated motility in the presence of VLY plus PMB (**Fig. 2D**) were similar to those observed with VLY alone.

**Figure 2:**
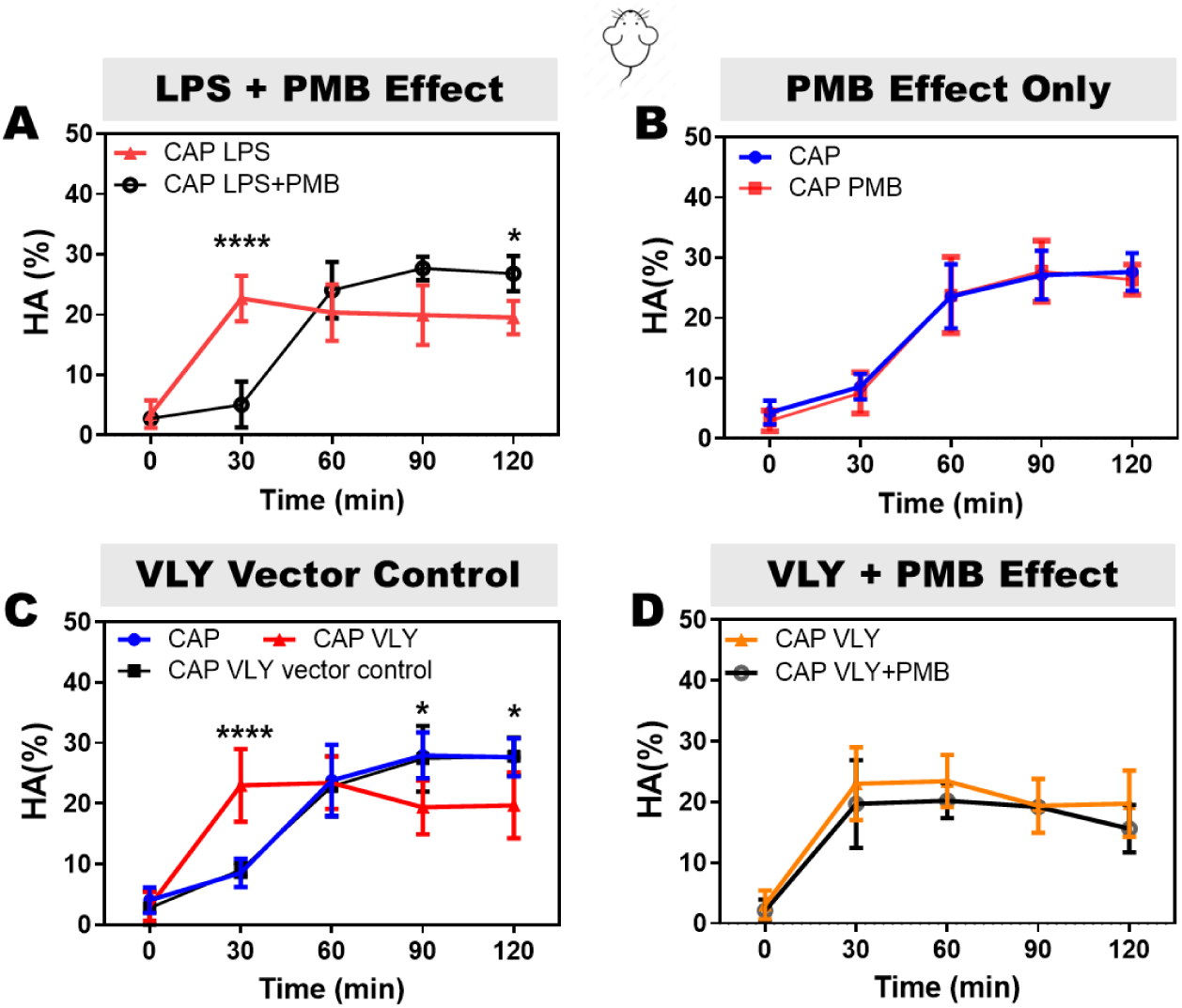
LPS and VLY effects on mouse sperm are specific. CASA measurements were obtained for hyperactivated motility of mouse sperm incubated under capacitating (CAP) conditions in the presence of **(A)** 1 µg/ml LPS or 100 µg/ml PMB + 1 µg/ml LPS, **(B)** vehicle or 100 µg/ml PMB alone, **(C)** vehicle, 1 µg/ml VLY, or VLY empty vector control, and **(D)** 1 µg/ml VLY or 100 µg/ml PMB + 1 µg/ml VLY. Here, 0 min is the time-point of BSA + sodium bicarbonate addition to initiate *in vitro* capacitation. Data are presented as mean and standard deviation (n≥4 biological replicates). *P<0.05, ****P<0.0001. Data were analyzed by two-way ANOVA with Bonferroni’s multiple comparison test.

Because capacitation media contains 15 mM HCO_3_^−^, we wanted to determine whether the effects could be explained by changes in osmolarity. Therefore, we increased the external medium osmolarity with 15 mM NaCl and found that neither LPS nor VLY induced hyperactivated motility in this condition (**Fig. S4**). We conclude that the effects on hyperactivation were due to changes in the capacitation signaling pathway induced by HCO3^−^ and not by changes in external osmolarity (Ferreira *et al*., 2021b).

### LPS and VLY interfere with human sperm capacitation

To determine whether LPS and VLY also impair human sperm function, we obtained leftover sperm samples from patients with normal semen parameters being treated at the Reproductive Endocrinology and Infertility Clinic at Washington University in Saint Louis. As the median concentration of LPS in patients with BV is approximately 3000 EU/mL (Aroutcheva *et al*., 2008), we used 1000 EU/mL LPS, corresponding to 0.1 µg/mL LPS, for human sperm. The physiologically relevant concentration of VLY is unknown, so we used the same concentration as LPS. Immediately after adding 0.1 µg/mL LPS or 0.1 µg/mL VLY to sperm in CAP conditions, the percentage of hyperactivated sperm was significantly greater than in untreated controls (**Fig. 3A, B**). However, by the end of the 180-min treatment period, human sperm treated with LPS or VLY showed significantly lower hyperactivated, total, and progressive motility than untreated sperm (**Fig. 3A–F**). LPS and VLY had no effect on human sperm hyperactivation or total or progressive motility in non-CAP conditions (**Fig. S5**). PMB prevented the dose-dependent increase in human sperm hyperactivation induced by LPS **(Fig. S6)**.

**Figure 3:**
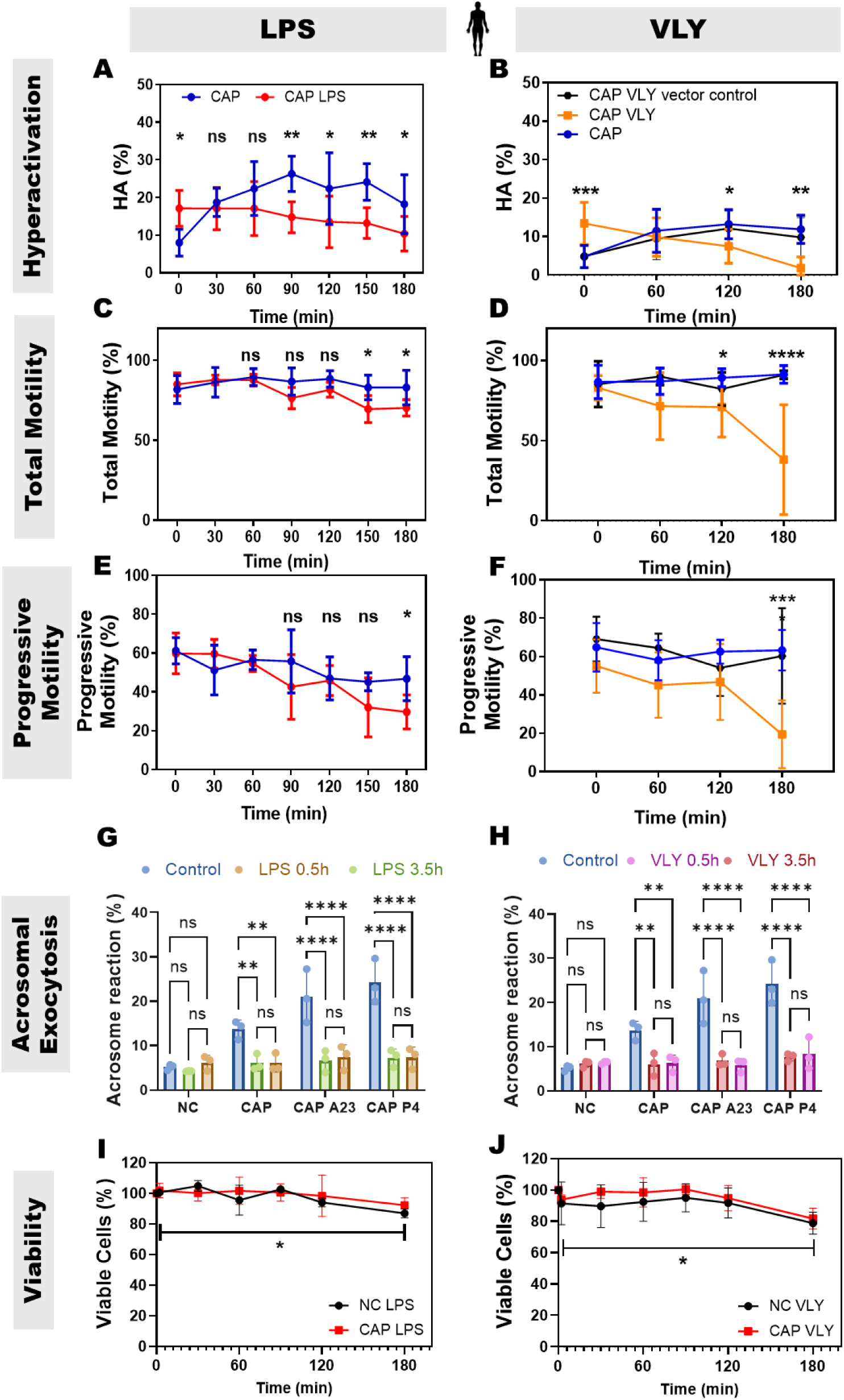
LPS and VLY impair human sperm hyperactivation, total motility, progressive motility, and acrosomal exocytosis in capacitating conditions. CASA measurements were obtained for (**A, B**) hyperactivated motility, (**C, D**) total motility, and (**E, F**) progressive motility of human sperm incubated under capacitating (CAP) conditions in the presence and absence of 0.1 µg/mL **(A, C, E)** LPS (n=5 biological replicates) and **(B, D, F)** VLY or VLY vector control (n≥5 biological replicates). Here 0 min is the time-point of BSA + sodium bicarbonate addition to initiate *in vitro* capacitation. Acrosomal exocytosis (AE) was quantified in non-CAP (NC), and CAP sperm (200 each, n=3 biological replicates) incubated with **(G)** LPS and **(H)** VLY for 0.5 and 3.5 h. Similarly, AE induced by 10 µM A23187 or progesterone (P4), was also quantified in CAP sperm. Sperm viability for NC and CAP sperm was analyzed in the presence of 0.1 µg/ml **(I)** LPS or **(J)** VLY (10,000 sperm each, n≥3 biological replicates). Data were analyzed by two-way ANOVA with Bonferroni’s multiple comparison test and are presented as mean and standard deviation. *P<0.05, **P<0.01, ***P<0.001, ****P<0.0001.

Next, we measured the effect of a ten-fold lower dose of LPS and VLY (0.1 µg/mL) on human sperm acrosomal exocytosis induced by addition of the Ca^2+^ ionophore A23187 or progesterone. As reported earlier (Baro Graf *et al*., 2020), about 15–30% of human sperm underwent induced acrosomal exocytosis in control conditions **(Fig. 3G, H)**. Both LPS **(Fig. 3G)** and VLY **(Fig. 3H)** significantly reduced the percentage of acrosomal exocytosis induced by A23187 or progesterone in CAP conditions but had no effect in non-CAP conditions. At 180 min of treatment, LPS **(Fig. 3I)** and VLY **(Fig. 3J)** significantly reduced human sperm viability by 13% and 21.2%, respectively, in non-CAP conditions. Under CAP conditions, LPS and VLY reduced the viability by 7.7% and 18.3%, respectively; however, this was not statistically significant. We conclude that LPS and VLY impair human sperm function in a similar manner as they impair mouse sperm function.

### LPS and VLY rapidly and irreversibly induce increases in intracellular Ca^2+^ concentration in CAP mouse and human sperm

Because both hyperactivated motility and acrosomal exocytosis are dependent on Ca^2+^ influx and increased intracellular Ca^2+^ concentration ([Ca^2+^]_I_), we wondered whether LPS and VLY affected sperm [Ca^2+^]_I_. To test this idea, we loaded mouse sperm with the cell-permeant Ca^2+^-sensitive fluorescent dye Fluo-4 AM. In control conditions, mouse sperm [Ca^2+^]_I_ reached maximum after addition of CAP media, with a time of initiation of the response (tau) of 164.2 ± 61.91. Addition of LPS and VLY increased [Ca^2+^]_I_ more rapidly, with a tau of 53.68 ± 36.36 and 20.01 ± 14.13, respectively **(Fig. S7-8)**. Also, the amplitude of the increase in [Ca^2+^]_I_ was significantly larger in the presence of LPS (**Fig. S8**). We next compared the LPS and VLY effects between non-CAP and CAP conditions. Between 10% and 40% of mouse sperm in non-CAP conditions and 50-100% of sperm in CAP conditions reached maximum [Ca^2+^]_I_ after addition of LPS or VLY (**Fig. 4A–F**). The amplitude of the Ca^2+^ response was greater and the tau was shorter in CAP than in non-CAP conditions (**Fig. 4G–J**) (see traces of LPS- and VLY- induced [Ca^2+^]_I_ response in **Fig. S9**). Experiments using human sperm produced similar results (**Fig. 5**, **Fig. S10).** Washout experiments indicated that the LPS- and VLY-induced Ca^2+^ responses were irreversible for both mouse and human sperm (**Fig. 6**). Therefore, we showed that both LPS and VLY induced rapid and irreversible increases in [Ca^2+^]_I_, and these increases were significantly larger in capacitated conditions than in non-capacitated conditions.

**Figure 4:**
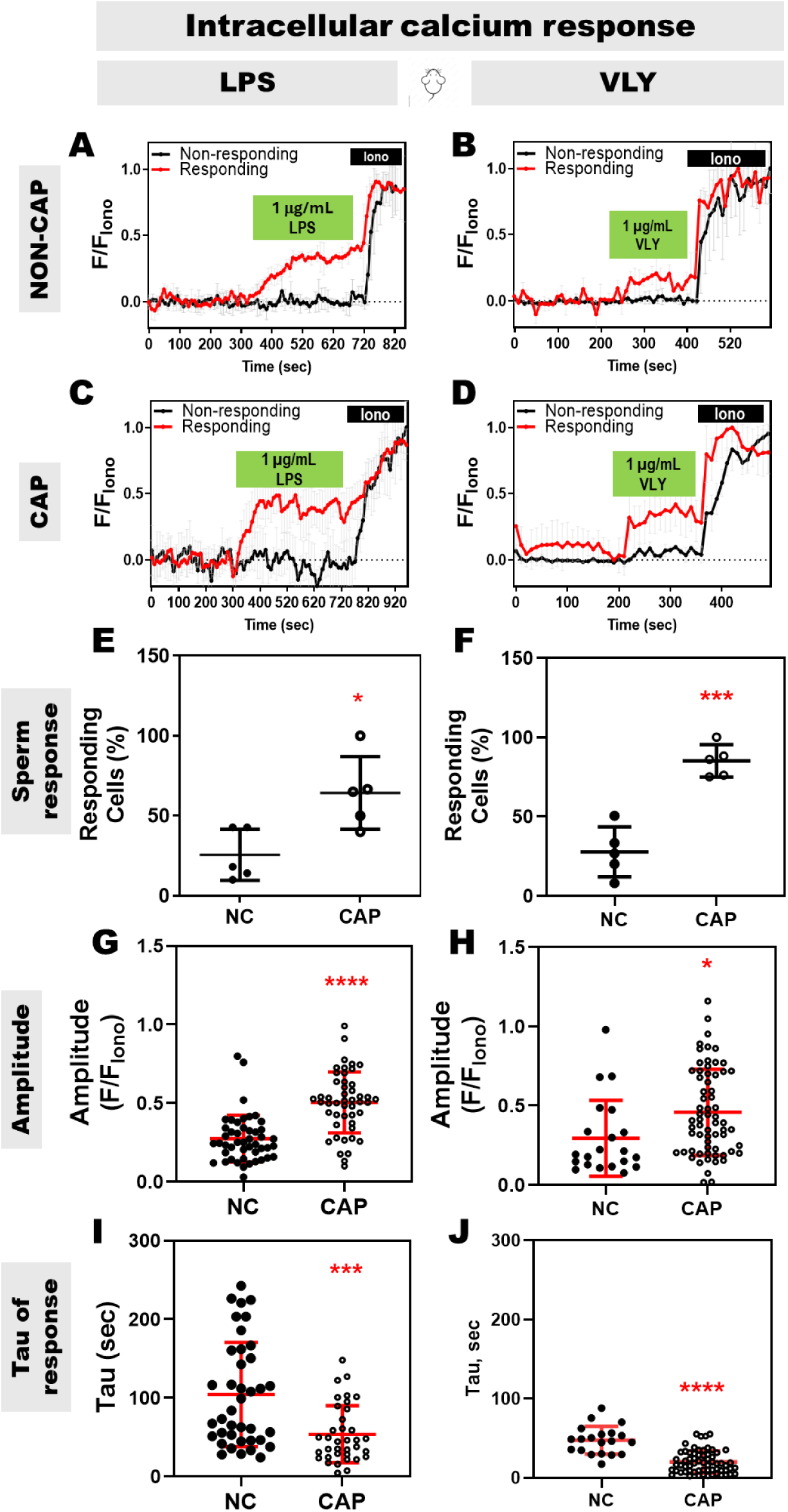
LPS and VLY cause rapid increases in [Ca^2+^]_I_ in capacitating mouse sperm. Representative traces of normalized Fluo4-AM fluorescence in responding (red) versus non-responding (black) mouse sperm with 1 µg/mL **(A, C)** LPS and **(B, D)** VLY, incubated under (**A, B**) Non-CAP (NC) and (**C, D**) CAP conditions. Each trace was normalized to its respective ionomycin (Iono, 2-5 µM) response. Percentages of mouse sperm responding to **(E)** LPS and **(F)** VLY, in NC and CAP conditions were calculated. (**G, H**) Amplitude and (**I, J**) tau of the [Ca^2+^]_I_ response in NC and CAP sperm, with 1 µg/mL **(G, I)** LPS and **(H, J)** VLY. Data are presented as mean and standard deviation (n=4 biological replicates). *P<0.05, ***P<0.001, ****P<0.0001, by unpaired t-test for **E-J**.

**Figure 5:**
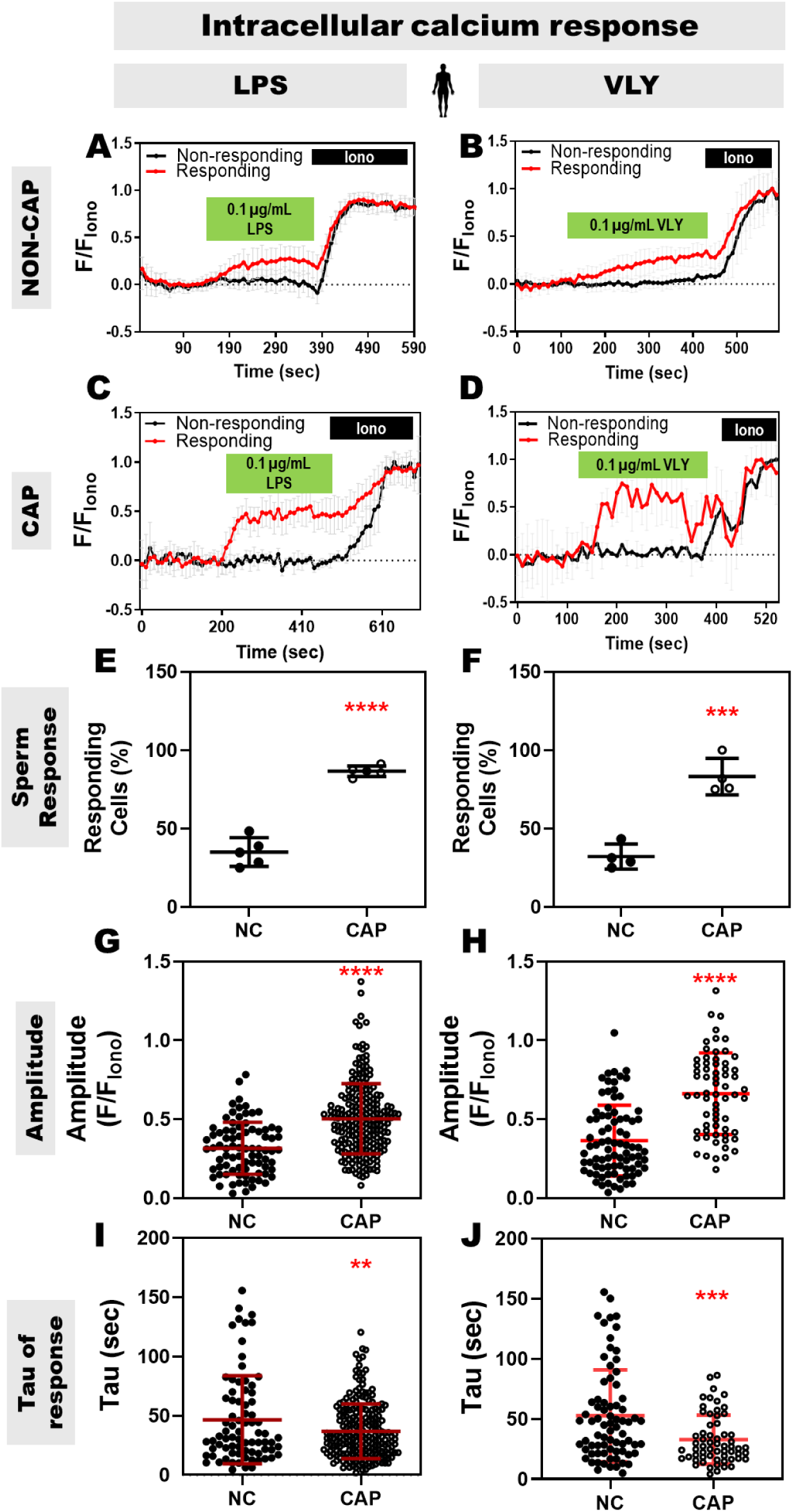
LPS and VLY cause rapid increases in [Ca^2+^]_I_ in capacitating human sperm. Representative traces of normalized Fluo4-AM fluorescence in responding (red) versus non-responding (black) human sperm with 0.1 µg/mL **(A, C)** LPS and **(B, D)** VLY, incubated under (**A, B**) Non-CAP (NC) and (**C, D**) CAP conditions. Each trace was normalized to its respective ionomycin (Iono, 2-5 µM) response. Percentages of human sperm responding to **(E)** LPS and **(F)** VLY, in NC and CAP conditions were calculated. (**G, H**) Amplitude and (**I, J**) tau of the [Ca^2+^]_I_ response in NC and CAP sperm, with 0.1 µg/mL **(G, I)** LPS, and **(H, J)** VLY was quantified. Data are presented as mean and standard deviation (n≥4 biological replicates). **P<0.01, ***P<0.001, ****P<0.0001, by unpaired t-test for **E-J**.

**Figure 6:**
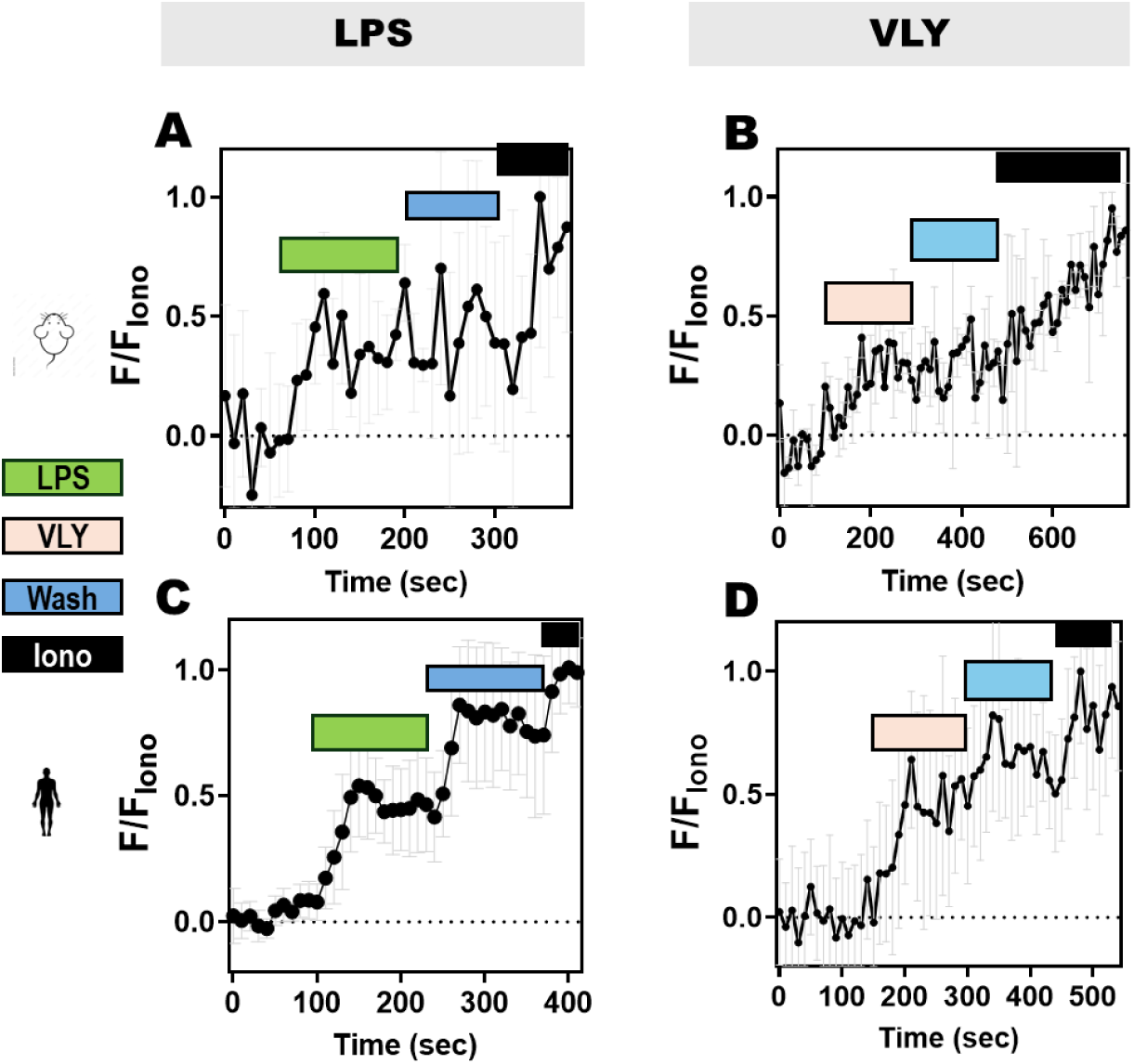
LPS- and VLY-induced [Ca^2+^]_I_ increases in mouse and human sperm are irreversible. Representative traces of normalized Fluo4-AM fluorescence in CAP (**A, B**) mouse and (**C, D**) human sperm, perfused with (**A, C**) LPS and (**B, D**) VLY, followed by media (Wash, denoted in blue), and subsequent perfusion with 2-5 µM ionomycin (Iono, denoted in black). Each trace was normalized to its respective ionomycin response. Data are presented as mean and standard deviation (n=3 biological replicates).

### LPS-induced Ca^2+^ response requires TLR4 and external Ca^2+^ but not CatSper

The main mechanism of [Ca^2+^]_I_ increase during normal capacitation is entry of external Ca^2+^through the sperm specific Ca^2+^ channel, CatSper. To determine whether the effects of LPS required external Ca^2+^, we exposed mouse sperm to CAP conditions in media that lacked Ca^2+^. In this condition, LPS did not cause a [Ca^2+^]_I_ increase (**Fig. 7A**). Similarly, LPS did not increase [Ca^2+^]_I_ in human sperm (**Fig. 7B**). To determine whether CatSper was required for the LPS-induced [Ca^2+^]_I_ increase, we measured [Ca^2+^]_I_ in sperm from wild-type (**Fig. 7C**) and CatSper knock-out (**Fig. 7D**) mice. Neither the percentages of sperm responding to LPS (**Fig. 7E**) nor the amplitude **(Fig. S11A)** or tau **(Fig. S11B)** of their responses differed significantly between sperm from wild-type and CatSper knock-out mice. Moreover, the [Ca^2+^]_I_ changes in LPS-responding cells were similar for wild-type (**Fig. 7F**) and CatSper knock-out (**Fig. 7G**) mouse sperm. These data indicate that LPS triggered an increase in [Ca^2+^]_I_ through a mechanism independent of CatSper.

**Figure 7:**
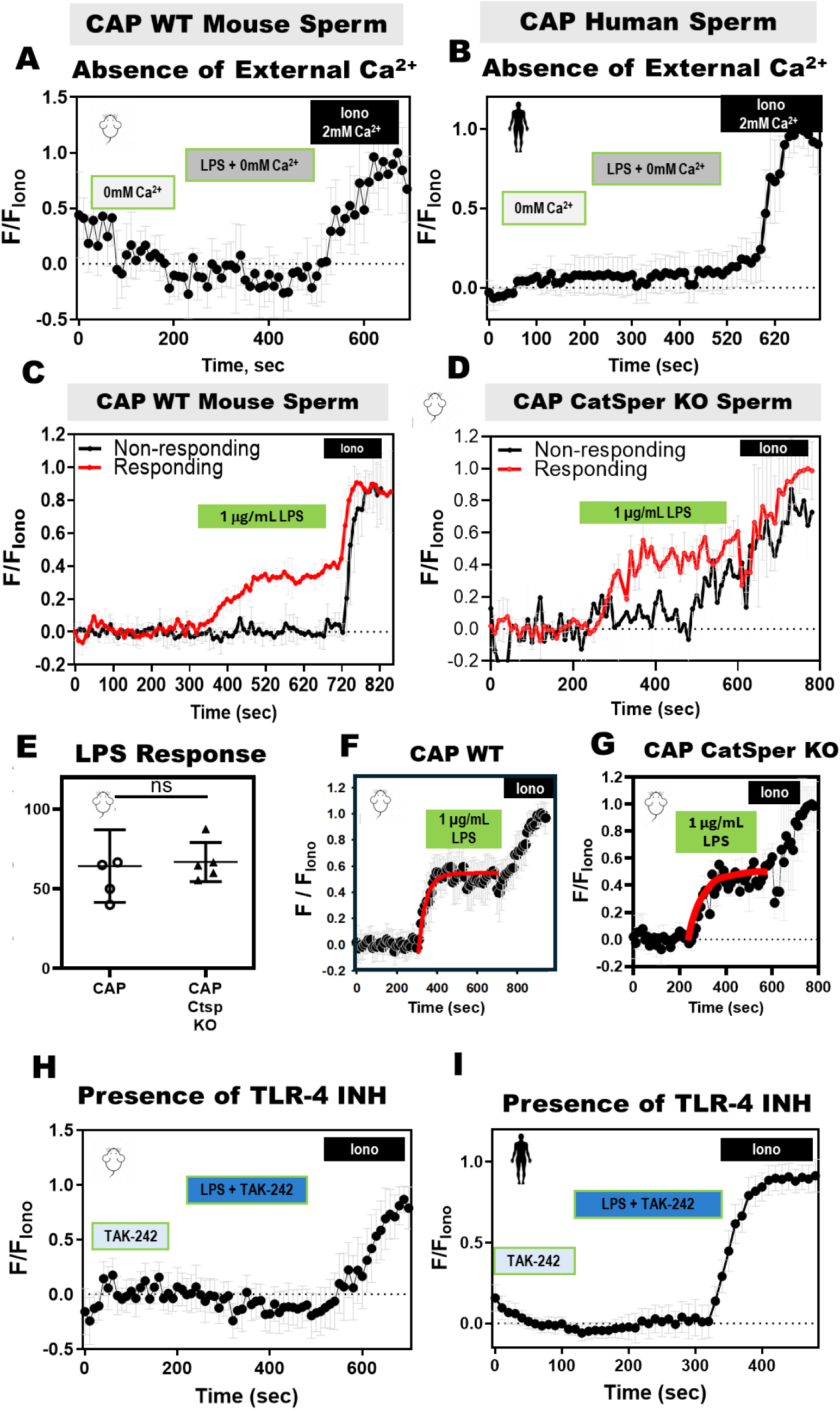
LPS-induced increase in sperm [Ca^2+^]_I_ is dependent on external Ca^2+^, independent of the CatSper ion channel, and mediated by the TLR4 receptor. Representative traces of normalized Fluo4-AM fluorescence from capacitating (CAP) **(A)** wild-type (WT) mouse, and **(B)** human sperm perfused with 0 mM Ca^2+^, 0 mM Ca^2+^ with LPS, and ionomycin (Iono) with 2 mM Ca^2+^ (n=3 biological replicates). Representative traces for LPS responding (red) and non-responding (black) **(C)** WT and **(D)** CatSper knock-out (KO) mouse sperm in the presence of 2 mM extracellular Ca^2+^. **(E)** Percentages of LPS-responding sperm with increased [Ca^2+^]_I_ from WT and CatSper KO mice (n=5 biological replicates). The LPS-induced [Ca^2+^]_I_ responses are shown on the traces obtained from **(F)** WT and **(G)** CatSper KO mouse sperm. Red curves calculated as a standard exponential fit. Representative traces of Fluo4-AM fluorescence from CAP **(H)** WT mouse, and **(I)** human sperm perfused with TAK-242 (TLR4 inhibitor), TAK-242 with LPS, and ionomycin (Iono), in the presence of 2 mM extracellular Ca^2+^ throughout the experiment (n=3 biological replicates). Each trace was normalized to its respective ionomycin (Iono, 2-5 µM) response. Data are presented as mean and standard deviation. ns= non-significant, by unpaired t-test in **(E).**

Given that LPS binding to the TLR4 receptor induces [Ca^2+^]_I_ increases in other cell types (Tauseef *et al*., 2012), we wondered whether TLR4 was required for LPS effects on [Ca^2+^]_I_ in sperm. To test this idea, we exposed sperm to LPS in the presence of the specific TLR4 inhibitor TAK-242 (Takashima *et al*., 2009; Yuko *et al*., 2020). The LPS-induced Ca^2+^ response did not occur in mouse (**Fig. 7H**) or human (**Fig. 7I**) sperm in the presence of TAK-242. These data show that the [Ca^2+^]_I_ increase induced by LPS in human and mouse sperm is mediated by the TLR4 receptor.

## Discussion

Bacterial vaginosis (BV) is more prevalent in infertile women than in fertile women, and 37.4% of females with unexplained infertility are diagnosed with BV (Salah *et al*., 2013). Although the cause of infertility among patients with BV is unclear, several mechanisms have been proposed. One possibility is that BV-related bacteria can induce immune activation and increase concentrations of proinflammatory cytokines, resulting in mucosal inflammation in the genital tract (van Teijlingen *et al*., 2020). Higher cervical concentrations of the cytokines interleukin (IL)-1β, IL-6, and IL-8 have been observed in women with infertility and BV (Spandorfer *et al*., 2001). Another possibility is that sialidases and other mucinases produced by microorganisms impair cervical mucus integrity (Wiggins *et al*., 2001) by degrading mucins. This could weaken the natural barrier against microbial invasion and facilitate bacterial adhesion and colonization, possibly leading to upper reproductive tract disease and infertility. Finally, women with BV have a 3.4 and 4.1-fold, respectively, elevated risk of acquiring the sexually transmitted infections with *Chlamydia trachomatis* and *Neisseria gonorrhoeae* (Wiesenfeld *et al*., 2003), which can impair fertility. Additionally, BV is associated with increased rates of upper genital tract infections and pelvic inflammatory disease, both of which are linked to infertility (Ravel *et al*., 2021).

Here, we present three lines of evidence supporting a new mechanism by which BV contributes to female infertility: the BV-associated toxins LPS and VLY interfere with sperm capacitation, which is required to fertilize an egg. First, both LPS and VLY treatment reduced the fertilizing ability of mouse sperm. Second, LPS and VLY impaired hyperactivation, motility, and acrosomal exocytosis in both mouse and human sperm. Third, LPS and VLY caused premature and irreversible increases in intracellular Ca^2+^ concentration ([Ca^2+^]_I_) in both mouse and human sperm.

Our results regarding the impact of LPS on mouse sperm are consistent with findings reported by Fujita *et. al.* (Fujita *et al*., 2011). They reported decreased mouse sperm motility, increased sperm apoptosis, and decreased fertilization ability after 6 h of incubation with LPS. We observed that incubation with LPS for over 90 min led to decreased sperm motility and a nearly 50% reduction in the formation of two pronuclei-stage embryos. We also noted a slight decrease in viability after 3 h in capacitated media. However, the largest effects we observed were on hyperactivated motility and acrosomal exocytosis, which were not explored by Fujita and co-workers.

Li *et. al*. also found that LPS decreased human sperm motility and intracellular cAMP concentrations (Li *et al*., 2016). Inconsistent with our findings, Li *et. al.* reported that LPS did not affect sperm [Ca^2+^]_I_, acrosomal exocytosis, or capacitation (Li *et al*., 2016). These negative results may have been due to contamination of controls or reagents with LPS. Alternatively, LPS can adhere to tubes upon dilution and be lost. We used methods to prevent these problems and found that the effects of LPS were irreversible. Given this irreversibility, we speculate that LPS in the vagina in patients with BV could affect sperm fertility. Consistent with our findings, Sahnoun *et. al.* reported that LPS led to reduced percentages of sperm acrosomal exocytosis in infertile males (Sahnoun *et al*., 2017). Additionally, LPS was reported to decrease human sperm motility and curvilinear movement, inhibit phosphorylation of NF-κB inhibitor alpha, and cause increased β-oxidation of fatty acids (Li *et al*., 2023).

Previous results regarding the effects of LPS on human sperm viability have been contradictory. In one report, 0.1 to 100 µg/mL LPS had no effect on viability (Li *et al*., 2016), whereas in another report, 50 µg/mL LPS caused 80% reduction in human sperm viability (Galdiero *et al*., 1988). With our validated concentrations of high-quality LPS and precautions to avoid loss of LPS through adsorption, we observed only a small decrease in sperm viability after 180 min of treatment. This small effect is unlikely to account for the effects of LPS on motility or acrosomal exocytosis.

Both sperm hyperactivation and acrosomal exocytosis are finely regulated by intracellular Ca^2+^. Calcium entry is required for both the initiation and maintenance of hyperactivation through direct regulation of axonemal machinery components (Carlson *et al*., 2003). However, excessive increases in [Ca^2+^]_I_ in mammalian sperm are associated with impaired fertilization potential. This includes mitochondrial failure, motility loss, activation of apoptotic cascades, and premature loss of the acrosome, known as spontaneous acrosome reaction (Liu and Baker, 1990). Prematurely inducing the human acrosome reaction with the Ca^2+^ ionophore A23187 decreases sperm-zona pellucida binding, causing fertilization failure *in vitro* (Liu and Baker, 1990). Additionally, spermatozoa deficient in the Ca^2+^ pump PMCA4 cannot extrude Ca^2+^ and thus experience Ca^2+^ overload, resulting in male infertility (Okunade *et al*., 2004). Furthermore, high concentrations (10–20 μM) of the Ca^2+^ ionophore A23187 initially increase the sperm flagellar beat, then immobilize sperm after 10 min. Concentrations of 5–10 μM A23187 rapidly immobilized sperm by elevating [Ca²⁺]_I_, whereas lower concentrations (0.5 and 1 μM) caused smaller [Ca²⁺]_I_ increases and resulted in hyperactivation. Notably, A23187 in Ca²⁺-free medium did not immobilize sperm.

These findings suggest that excessive Ca²⁺ influx affects motility and, at high concentrations, can lead to sperm immobilization. Observing faster and higher-amplitude [Ca²⁺]_I_ increases in sperm exposed to LPS and VLY compared to controls, we speculate that both toxins cause sperm Ca^2+^ overload. However, the mechanisms by which this overload occurs in response to LPS and VLY might be distinct.

We found that LPS caused Ca^2+^ influx in both human and mouse sperm. In mammalian sperm, the Ca^2+^ influx into the flagellum is primarily regulated by CatSper (Kirichok *et al*., 2006), but we found that LPS-induced Ca^2+^ influx in mouse sperm was independent of CatSper. Fujita *et. al.* reported that the negative effects of LPS on sperm motility and fertility were blocked in TLR4 knockout mice (Fujita *et al*., 2011), suggesting that TLR4 participates in LPS-mediated Ca^2+^ signalling. Our experiments with the TLR4 inhibitor support the idea that the LPS-induced [Ca^2+^]_I_ increase in human and mouse sperm was mediated by the TLR4 receptor. Future experiments will be conducted to define the mechanisms by which this occurs.

VLY is a cytolysin that forms pores in the cell membrane (Ragaliauskas *et al*., 2019). In addition to causing cells to leak their intracellular contents (Randis *et al*., 2013; Morrill *et al*., 2023) the disruptions to the membrane would likely also allow extracellular Ca^2+^ to enter the cytoplasm, as we saw in both mouse and human sperm. Because VLY embeds in the human cell membrane, it is expected to be irreversible. Thus, it was not surprising that we were unable to wash out the effects of VLY in sperm.

In conclusion, our *in vitro* results show that the BV-associated toxins LPS and VLY modulate sperm Ca^2+^ entry, leading to dysregulated hyperactivation, inhibition of acrosomal exocytosis, and impaired fertility. If these effects occur *in vivo*, then BV toxins in the vagina could impair key events necessary to prepare sperm for fertilization. This mechanism may contribute to infertility in patients with BV.

## Author’s role

S.B. was involved in study design, data collection, analysis, and manuscript preparation. C.S., A.L., and W.L. provided their expertise and were responsible for the study design, conceptualization, supervision, and review of the manuscript. S.B., L.A., R.M., J.F., C.S., A.L., and W.L. contributed to the methodology, data collection, analysis, and investigation. Data visualization was performed by S.B., P.L., E.L., S.S, and A.G. Initial standardisation on VLY cloning and expression was done by V.R. LPS testing of reagents was done by C.A. VLY protein was prepared by S.M. and H.Z. All authors reviewed and contributed to the final manuscript.

## Acknowledgements

The authors are grateful to Deborah J. Frank for editing the manuscript, and Dr. Ali Ahmady, Lab Director of Reproductive Endocrinology and Infertility at Washington University in Saint Louis for facilitating the collection of human samples.

## Funding

This work was supported by the National Institutes of Health (grant #R01 HD069631 to CS).

## Conflict of interest

The authors declare that they have no conflict of interest.

## Data and materials availability

All data are available in the main text or the supplementary materials.

## Notes

### Competing Interest Statement

The authors have declared no competing interest.

## References

Allsworth JE, Peipert JF. Prevalence of Bacterial Vaginosis. Obstet Gynecol 2007;109:114– 120.

Aroutcheva A, Zaodung L, Faro S. Prevotella bivia as a Source of Lipopolysaccharide in the Vagina. Anaerobe 2008;14:256–260.

Baro Graf C, Ritagliati C, Torres-Monserrat V, Stival C, Carizza C, Buffone MG, Krapf D. Membrane Potential Assessment by Fluorimetry as a Predictor Tool of Human Sperm Fertilizing Capacity. Front Cell Dev Biol 2020;7:1–10.

Campisciano G, Iebba V, Zito G, Luppi S, Martinelli M, Fischer L, Seta F De, Basile G, Ricci G, Comar M. Lactobacillus iners and gasseri, prevotella bivia and hpv belong to the microbiological signature negatively affecting human reproduction. Microorganisms 2021;9:1–12.

Carlson AE, Westenbroek RE, Quill T, Ren D, Clapham DE, Hille B, Garbers DL, Babcock DF. CatSper1 required for evoked Ca2+ entry and control of flagellar function in sperm. PNAS 2003;100:14864–14868.

Chávez JC, Ferreira JJ, Butler A, La Vega Beltrán JL De, Treviño CL, Darszon A, Salkoff L, Santi CM. SLO3 K+ channels control calcium entry through CATSPER channels in sperm. J Biol Chem 2014;**289**:32266–32275.

Cocomazzi G, Stefani S De, Pup L Del, Palini S, Buccheri M, Primiterra M, Sciannamè N, Faioli R, Maglione A, Baldini GM, et al. The Impact of the Female Genital Microbiota on the Outcome of Assisted Reproduction Treatments. Microorganisms 2023;11:.

Criteria D, Associations E. Nonspecific Vaginitis Diagnostic Criteria and Microbial and Epidemiologic Associations. 1983;74:.

Eckert LO, Moore DE, Patton DL, Agnew KJ, Eschenbach DA. Relationship of vaginal bacteria and inflammation with conception and early pregnancy loss following in-vitro fertilization. Infect Dis Obs Gynecol 2003;11:11–17.

Ferreira JJ, Cassina A, Irigoyen P, Ford M, Pietroroia S, Peramsetty N, Radi R, Santi CM, Sapiro R. Increased mitochondrial activity upon CatSper channel activation is required for mouse sperm capacitation. Redox Biol 2021a;48:102176 (1-11). Elsevier B.V.

Ferreira JJ, Lybaert P, Puga-Molina LC, Santi CM. Conserved Mechanism of Bicarbonate-Induced Sensitization of CatSper Channels in Human and Mouse Sperm. Front Cell Dev Biol 2021b;9:1–14.

Firmal P, Shah VK, Chattopadhyay S. Insight Into TLR4-Mediated Immunomodulation in Normal Pregnancy and Related Disorders. Front Immunol 2020;11:1–16.

Fujita Y, Mihara T, Okazaki T, Shitanaka M, Kushino R, Ikeda C, Negishi H, Liu Z, Richards JS, Shimada M. Toll-like receptors (TLR) 2 and 4 on human sperm recognize bacterial endotoxins and mediate apoptosis. Hum Reprod 2011;26:2799–2806.

Galdiero F, Gorga F, Bentivoglio C, Mancuso R, Galdiero E, Tufano MA. The action of LPS porins and peptidoglycan fragments on human spermatozoa. Infection 1988;16:349–353.

George SD, Gerwen OT Van, Dong C, Sousa LG V, Cerca N, Elnaggar JH, Taylor CM, Muzny CA. The Role of Prevotella Species in Female Genital Tract Infections. Pathogens 2024;13:1–14.

Haahr T, Zacho J, Brauner M, Shathmigha K, Skov Jensen J, Humaidan P. Reproductive outcome of patients undergoing in vitro fertilisation treatment and diagnosed with bacterial vaginosis or abnormal vaginal microbiota : a systematic PRISMA review and meta-analysis. BJOG 2018;200–207.

Hasuwa H, Muro Y, Ikawa M, Kato N, Tsujimoto Y, Okabe M. Transgenic mouse sperm that have green acrosome and red mitochondria allow visualization of sperm and their acrosome reaction in vivo. Exp Anim 2010;59:105–107.

Hong X, Ma J, Yin J, Fang S, Geng J, Zhao H, Zhu M, Ye M, Zhu X, Xuan Y, et al. The association between vaginal microbiota and female infertility : a systematic review and meta - analysis. Arch Gynecol Obstet [Internet] 2020;302:569–578. Springer Berlin Heidelberg.

Kirichok Y, Navarro B, Clapham DE. Whole-cell patch-clamp measurements of spermatozoa reveal an alkaline-activated Ca2+ channel. Nature 2006;439:737–740.

La Vega-Beltran JL De, Sánchez-Cárdenas C, Krapf D, Hernandez-González EO, Wertheimer E, Treviño CL, Visconti PE, Darszon A. Mouse sperm membrane potential hyperpolarization is necessary and sufficient to prepare sperm for the acrosome reaction. J Biol Chem 2012;287:44384–44393.

Łaniewski P, Herbst-Kralovetz MM. Bacterial vaginosis and health-associated bacteria modulate the immunometabolic landscape in 3D model of human cervix. Biofilms and microbiomes 2021;7:1–17.

Li Y, Hu Y, Wang Z, Lu T, Yang Y, Diao H, Zheng X, Xie C, Zhang P, Zhang X, et al. IKBA phosphorylation governs human sperm motility through ACC-mediated fatty acid beta-oxidation. Commun Biol 2023;6:1–12. Springer US.

Li Z, Zhang D, He Y, Ding Z, Mao F, Zhang X. Lipopolysaccharide compromises human sperm function by reducing intracellular cAMP. Tohoku J Exp Med 2016;238:105–112.

Liu DY, Baker HWG. Inducing the human acrosome reaction with a calcium ionophore A23187 decreases sperm-zona pellucida binding with oocytes that failed to fertilize in vitro. J Reprod Fertil 1990;89:127–134.

Lyon M, Li P, Ferreira J, Lazarenko R, Kharade S, Kramer M, McClenahan S, Days E, Bauer J, Spitznagel B, et al. A selective inhibitor of the sperm-specific potassium channel SLO3 impairs human sperm function. Proc Natl Acad Sci 2023;120:e2212338120 (1-9).

Magata F, Tsukamura H, Matsuda F. The impact of inflammatory stress on hypothalamic kisspeptin neurons : Mechanisms underlying inflammation-associated infertility in humans and domestic animals. Peptides 2023;162:170958. Elsevier Inc.

Mania-pramanik J, Kerkar ÃSC, Mehta PB, Potdar S, Salvi VS. Use of Vaginal pH in Diagnosis of Infections and Its Association With Reproductive Manifestations. J Clin Lab Anal 2008;22:375–379.

Molina LCP, Gunderson S, Riley J, Lybaert P, Borrego-Alvarez A, Jungheim ES, Santi CM. Membrane Potential Determined by Flow Cytometry Predicts Fertilizing Ability of Human Sperm. Front Cell Dev Biol 2020;7:1–12.

Molina LCP, Luque GM, Balestrini PA, Marín-Briggiler CI, Romarowski A, Buffone MG. Molecular basis of human sperm capacitation. Front Cell Dev Biol 2018;6:1–23.

Morrill SR, Saha S, Varki AP, Lewis WG, Ram S, Lewis AL. Gardnerella Vaginolysin Potentiates Glycan Molecular Mimicry by Neisseria gonorrhoeae. J Infect Dis 2023;**228**:1610–1620. Oxford University Press.

Mortimer ST, Swan MA, Mortimer D. Effect of seminal plasma on capacitation and hyperactivation in human spermatozoa. Hum Reprod 1998;13:2139–2146.

Murphy K, Mitchell CM. The Interplay of Host Immunity , Environment and the Risk of Bacterial Vaginosis and Associated Reproductive Health Outcomes. J Infect Dis 2016;214:29–35.

Nakagata N. “Cryopreservation of Mouse Spermatozoa and In Vitro Fertilization” in Methods in Molecular Biology, Chapter 4, Pgs. 57-73. Methods Mol Biol Chapter *4* 2010;

O’Doherty AM, Fenza M Di, Kölle S. Lipopolysaccharide (LPS) disrupts particle transport, cilia function and sperm motility in an ex vivo oviduct model. Sci Rep 2016;**6**:1–11. Nature Publishing Group.

Okunade GW, Miller ML, Pyne GJ, Sutliff RL, O’Connor KT, Neumann JC, Andringa A, Miller DA, Prasad V, Doetschman T, et al. Targeted ablation of plasma membrane Ca2+-ATPase (PMCA) 1 and 4 indicates a major housekeeping function for PMCA1 and a critical role in hyperactivated sperm motility and male fertility for PMCA4. J Biol Chem 2004;279:33742–33750.

Oostrum N Van, Sutter P De, Meys J, Verstraelen H. Risks associated with bacterial vaginosis in infertility patients : a systematic review and meta-analysis. Hum Reprod 2013;28:1809–1815.

Ragaliauskas T, Plečkaitytė M, Jankunec M, Labanauskas L, Baranauskiene L, Valincius G. Inerolysin and vaginolysin, the cytolysins implicated in vaginal dysbiosis, differently impair molecular integrity of phospholipid membranes. Sci Rep 2019;9:1–11.

Randis TM, Kulkarni R, Aguilar JL, Ratner AJ. Antibody-based detection and inhibition of vaginolysin, the Gardnerella vaginalis cytolysin. PLoS One 2009;4:1–5.

Randis TM, Zaklama J, LaRocca TJ, Los FCO, Lewis EL, Desai P, Rampersaud R, Amaral FE, Ratner AJ. Vaginolysin drives epithelial ultrastructural responses to gardnerella vaginalis. Infect Immun 2013;81:4544–4550.

Ravel J, Gajer P, Abdo Z, Schneider GM, Koenig SSK, McCulle SL, Karlebach S, Gorle R, Russell J, Tacket CO, et al. Vaginal microbiome of reproductive-age women. Proc Natl Acad Sci U S A 2011;108:4680–4687.

Ravel J, Moreno I, Simón C. Bacterial vaginosis and its association with infertility, endometritis, and pelvic inflammatory disease. Am J Obstet Gynecol 2021;224:251–257. Elsevier Inc.

Sahnoun S, Sellami A, Chakroun N, Mseddi M, Attia H, Rebai T, Lassoued S. Human sperm Toll-like receptor 4 (TLR4) mediates acrosome reaction, oxidative stress markers, and sperm parameters in response to bacterial lipopolysaccharide in infertile men. J Assist Reprod Genet 2017;34:1067–1077. Journal of Assisted Reproduction and Genetics.

Salah RM, Allam AM, Magdy AM, Mohamed AS. Bacterial vaginosis and infertility: Cause or association? Eur J Obstet Gynecol Reprod Biol 2013;167:59–63. Elsevier Ireland Ltd.

Salminen A, Paananen R, Vuolteenaho R, Metsola J, Ojaniemi M, Autio-Harmainen H, Hallman M. Maternal endotoxin-induced preterm birth in mice: Fetal responses in toll-like receptors, collectins, and cytokines. Pediatr Res 2008;63:280–286.

Sosa CM, Pavarotti MA, Zanetti MN, Zoppino FCM, Blas GA De, Mayorga LS. Kinetics of human sperm acrosomal exocytosis. Mol Hum Reprod 2014;21:244–254.

Spandorfer SD, Neuer A, Paulo GC, Rosenwaks Z, Witkin S. Relationship of abnormal vaginal flora, proinflammatory cytokines and idiopathic infertility in women undergoing IVF. J Reprod Med 2001;46:806–810.

Suarez SS. Control of hyperactivation in sperm. Hum Reprod Update 2008;14:647–657.

Takashima K, Matsunaga N, Yoshimatsu M, Hazeki K, Kaisho T, Uekata M, Hazeki O, Akira S, Iizawa Y, Ii M. Analysis of binding site for the novel small-molecule TLR4 signal transduction inhibitor TAK-242 and its therapeutic effect on mouse sepsis model Abbreviations : Br J Pharmacol 2009;**157**:1250–1262.

Tauseef M, Knezevic N, Chava KR, Smith M, Sukriti S, Gianaris N, Obukhov AG, Vogel SM, Schraufnage DE, Dietrich A, et al. TLR4 activation of TRPC6-dependent calcium signaling mediates endotoxininduced lung vascular permeability and inflammation. J Exp Med 2012;209:1953–1968.

Teijlingen NH van, Helgers LC, Zijlstra - Willems EM, Hamme JL van, Ribeiro CMS, Strijbis K, Geijtenbeek TBH. Vaginal dysbiosis associated-bacteria Megasphaera elsdenii and Prevotella timonensis induce immune activation via dendritic cells. J Reprod Immunol 2020;138:103085 (1-8). Elsevier.

Turpin R, Tuddenham S, He X, Klebanoff MA, Ghanem KG, Brotman RM. Bacterial Vaginosis and Behavioral Factors Associated with Incident Pelvic Inflammatory Disease in the Longitudinal Study of Vaginal Flora. J Infect Dis 2021;224:S137–S144.

Wang X, Rousset CI, Hagberg H, Mallard C. Lipopolysaccharide-induced inflammation and perinatal brain injury. Semin Fetal Neonatal Med 2006;11:343–353.

Wiesenfeld HC, Hillier SL, Krohn MA, Landers D V, Sweet RL. Bacterial Vaginosis Is a Strong Predictor of Neisseria gonorrhoeae and Chlamydia trachomatis Infection. Clin Infect Dis 2003;36:663–668.

Wiggins R, Hicks SJ, Soothill PW, Millar MR, Corfield AP. Mucinases and sialidases: Their role in the pathogenesis of sexually transmitted infections in the female genital tract. Sex Transm Infect 2001;77:402–408.

Yoo DK, Lee S-H. Effect of Lipopolysaccharide (LPS) Exposure on the Reproductive Organs of Immature Female Rats. Dev Reprod 2016;20:113–121.

Yuko O, Yuko M, Masafumi S, Kazuho S, Shoichiro H, Shimomura K, Inoue S, Kotani J. TAK-242 , a specific inhibitor of Toll-like receptor 4 signalling , prevents endotoxemia-induced skeletal muscle wasting in mice. Sci Rep 2020;10:1–13.

