## Supplementary figures for "Bacterial Vaginosis Toxins Impair Sperm Capacitation and Fertilization"

### Fig. S1

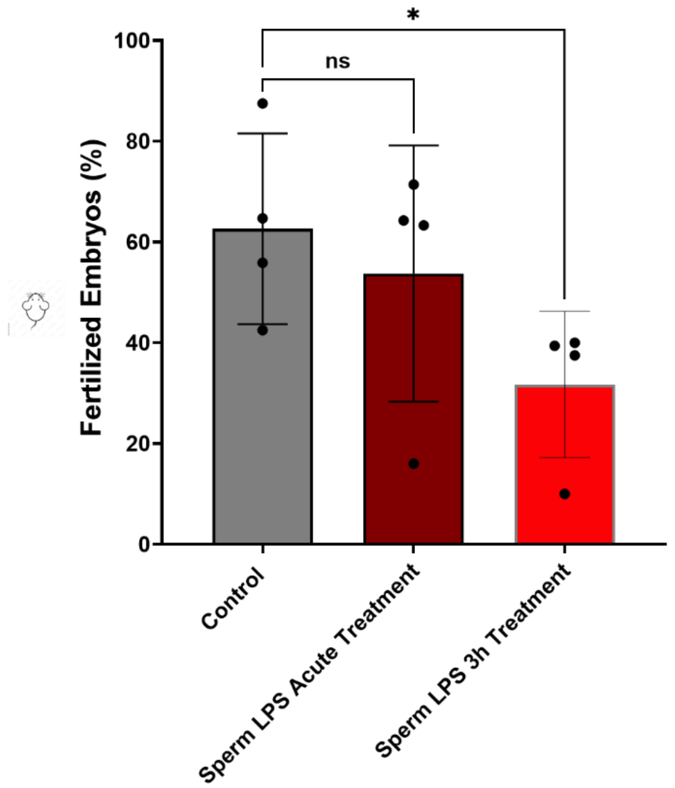


**Fig. S1. Mouse capacitating sperm fertilization rates decrease after prolonged treatment with LPS but not with acute exposure.** Percentage of oocytes that were fertilized in vitro and reached the 2-cell stage embryo after 24 h. Sperm were treated over 3 h in the absence (Control) or presence of 1 µg/mL LPS (sperm LPS 3 h treatment) or were only exposed to LPS in the fertilization drop (sperm LPS acute treatment) under capacitating conditions. Data are presented as mean and standard deviation (n=4 biological replicates). *P<0.05 by one-way ANOVA was performed with Bonferroni's multiple comparison test.

**Fig. S2**

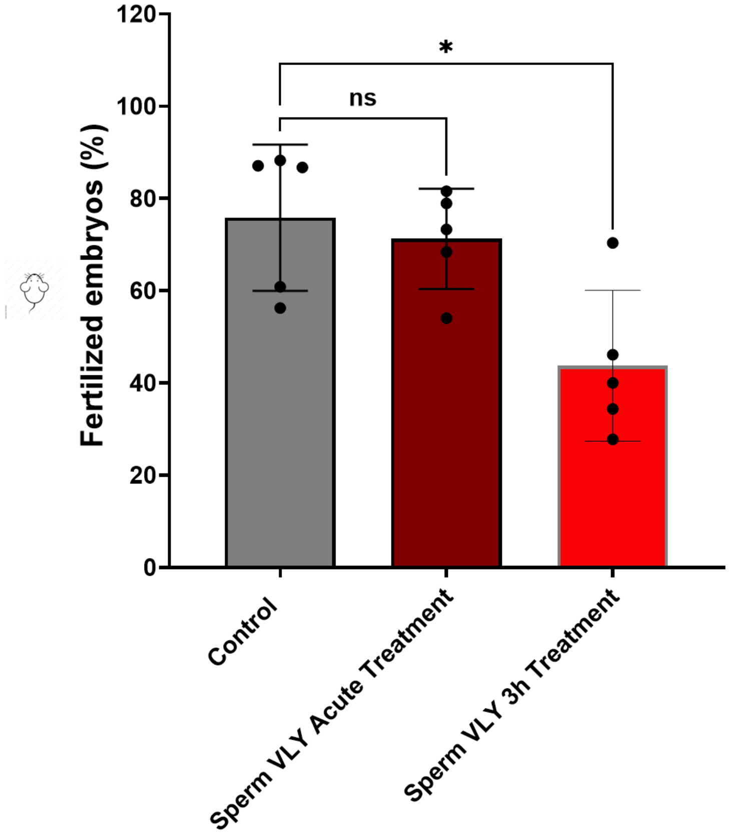


**Fig. S2. Capacitating mouse sperm fertilization rates decrease after prolonged treatment with VLY but not with acute exposure.** Percentage of oocytes that were fertilized *in vitro* and reached the 2-cell stage embryo after 24 h. Sperm were treated over 3 h in the absence (Control) or presence of 1 µg/mL VLY (sperm VLY 3 h treatment) or were only exposed to VLY in the fertilization drop (sperm VLY acute treatment) under capacitating conditions. Data are presented as mean and standard deviation (n=5 biological replicates). *P<0.05 by one-way ANOVA with Bonferroni's multiple comparison test.

### Fig. S3

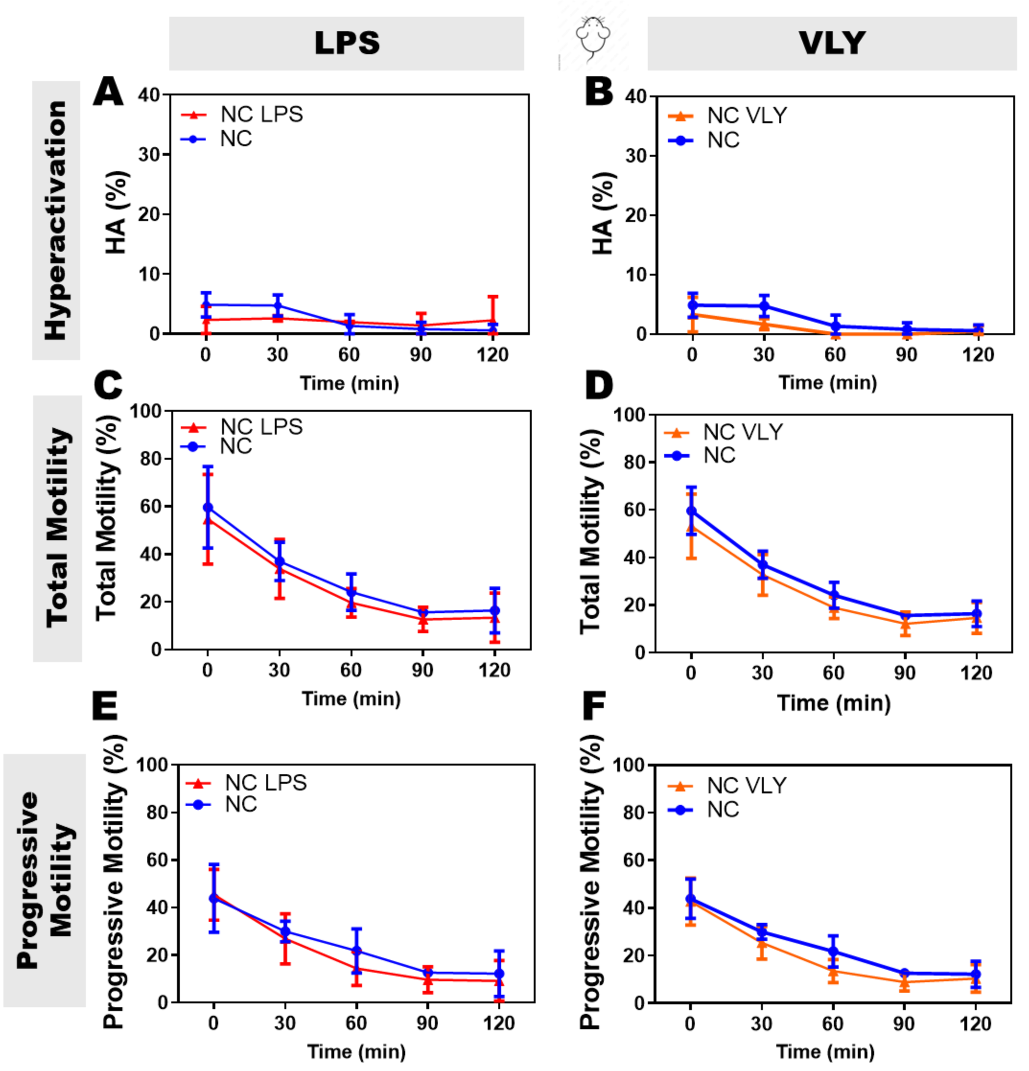


**Fig. S3. In non-capacitating conditions LPS and VLY do not affect mouse sperm hyperactivated, total, or progressive motility.** CASA measurements of **(A, B)** hyperactivated, **(C, D)** total, and **(E, F)** progressive motility of mouse sperm incubated under non-capacitating conditions, in the presence and absence of 1 µg/mL **(A, C, E)** LPS and **(B, D, F)** VLY, where 0 min is the time-point of BSA and sodium bicarbonate addition to the sperm suspension. Data are presented as mean and standard deviation (n=3 biological replicates).

#### Fig. S4

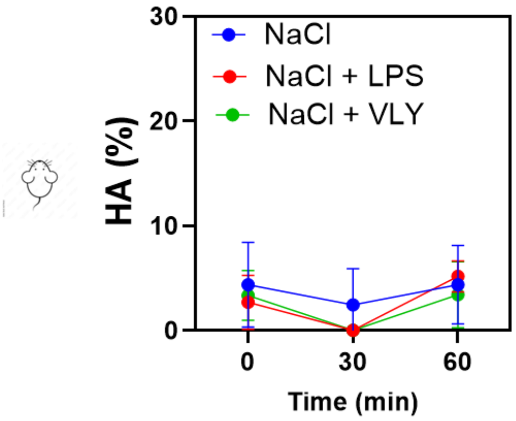


**Fig. S4. Increase in osmolarity by addition of sodium chloride into the external medium does not cause LPS- and VLY-induced mouse sperm hyperactivation.** The percentage of sperm with hyperactivated motility during capacitation, measured by CASA from sperm samples incubated in the presence and absence of 1 µg/mL LPS or VLY, where 0 min is the time point when 15 mM sodium chloride (NaCl), was added to the sperm suspension. Data are presented as mean and standard deviation (n=3 biological replicates). Data were evaluated by two-way ANOVA with Bonferroni's multiple comparison test.

**Fig. S5**

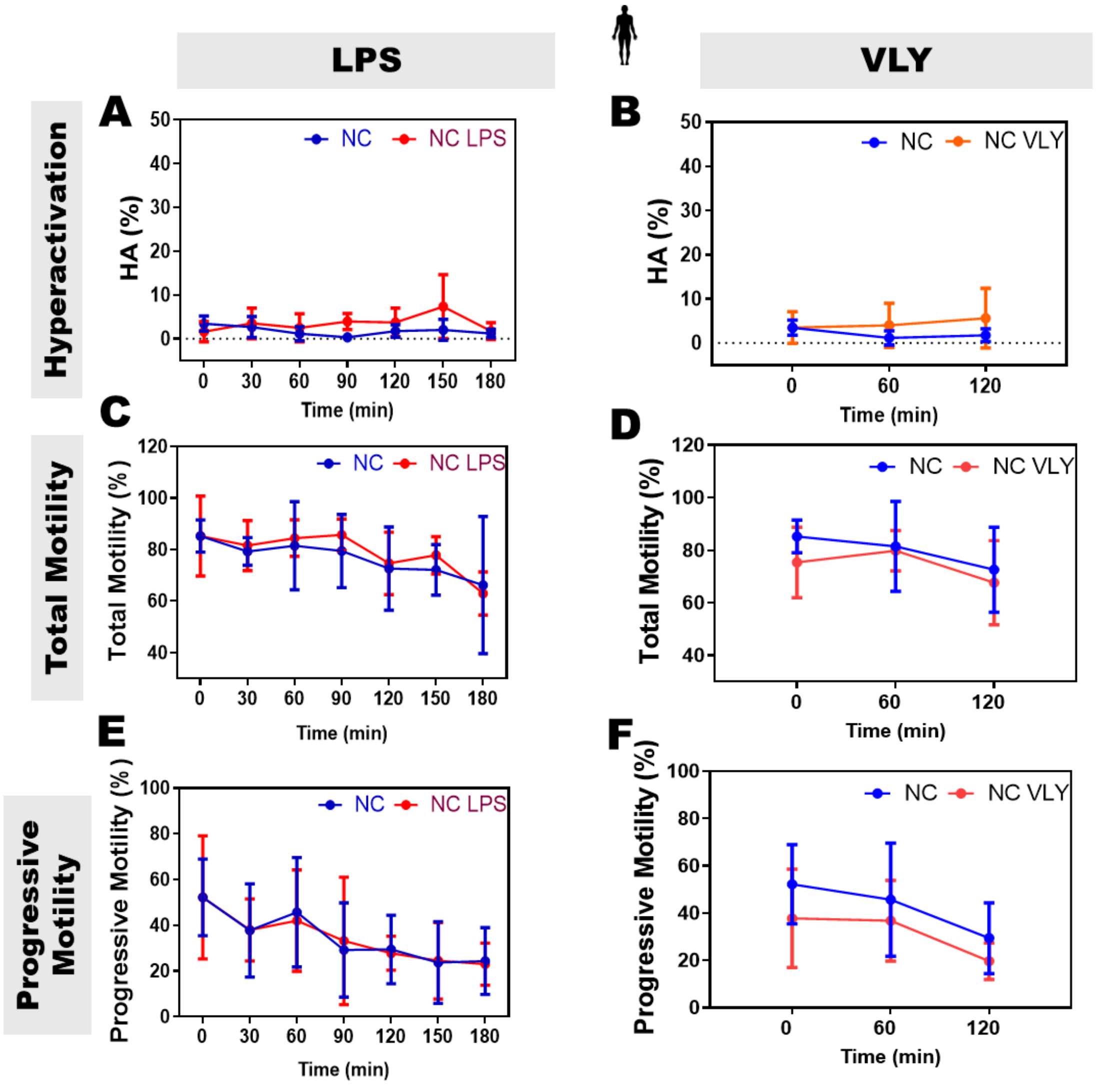


**Fig. S5. In non-capacitating conditions, LPS and VLY do not affect human sperm hyperactivated, total, or progressive motility.** CASA measurements of (**A, B**) hyperactivated, (**C, D**) total, and (**E, F**) progressive motility of human sperm in non-capacitating conditions in the presence and absence of 0.1 µg/mL **(A, C, E)** LPS and **(B, D, F)** VLY, where 0 min is the time-point of BSA and sodium bicarbonate addition to the sperm suspension. Data are presented as mean and standard deviation (n=5 biological replicates).

**Fig. S6**

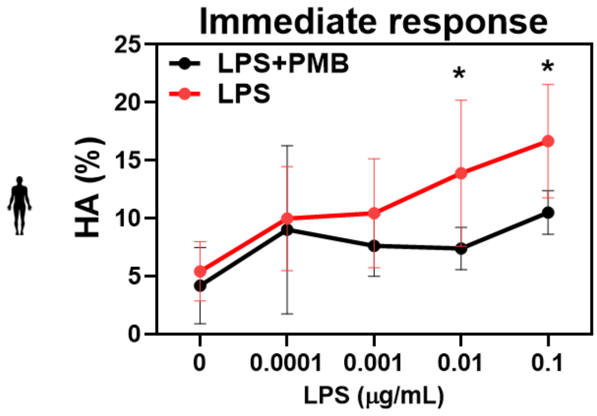


**Fig. S6. LPS-induced human sperm hyperactivation is dose-dependent and inhibited by polymyxin B:** The percentage of sperm with hyperactivated motility at time 0 of capacitation measured by CASA in sperm samples incubated in the presence of increasing concentrations of LPS with or without 100 µg/ml PMB. Data are presented as mean and standard deviation (n=5 biological replicates). *P<0.05.

#### Fig. S7

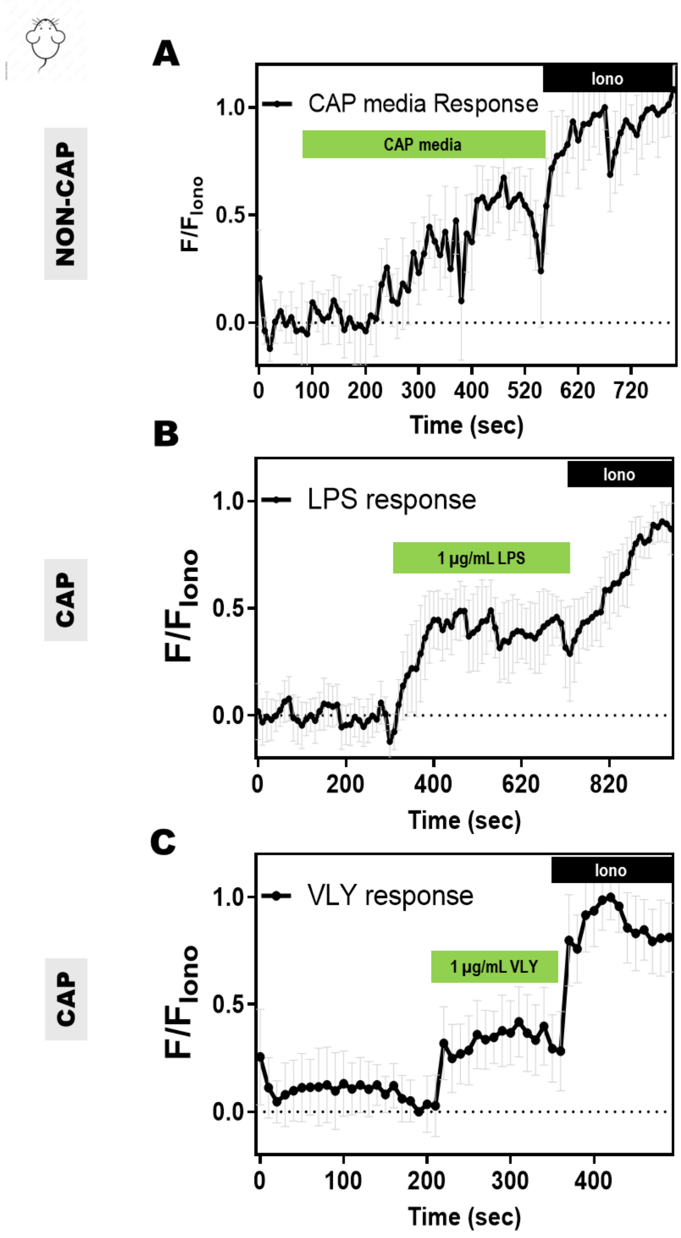


**Fig. S7. Sperm [Ca$^{2+}$]$_I$ response in WT mouse sperm is faster in the presence than in the absence of LPS or VLY.** Representative traces of [Ca$^{2+}$]$_I$ response in **(A)** Non-Capacitating (NON-CAP) mouse sperm when exposed to capacitating (CAP) media, followed by 2-5 µM Ionomycin (Iono). Similar traces were obtained for capacitating (CAP) mouse sperm when exposed to **(B)** LPS and **(C)** VLY. Each trace was normalized to its respective ionomycin (Iono) response. Data are presented as mean and standard deviation (n=3 biological replicates).

#### Fig. S8

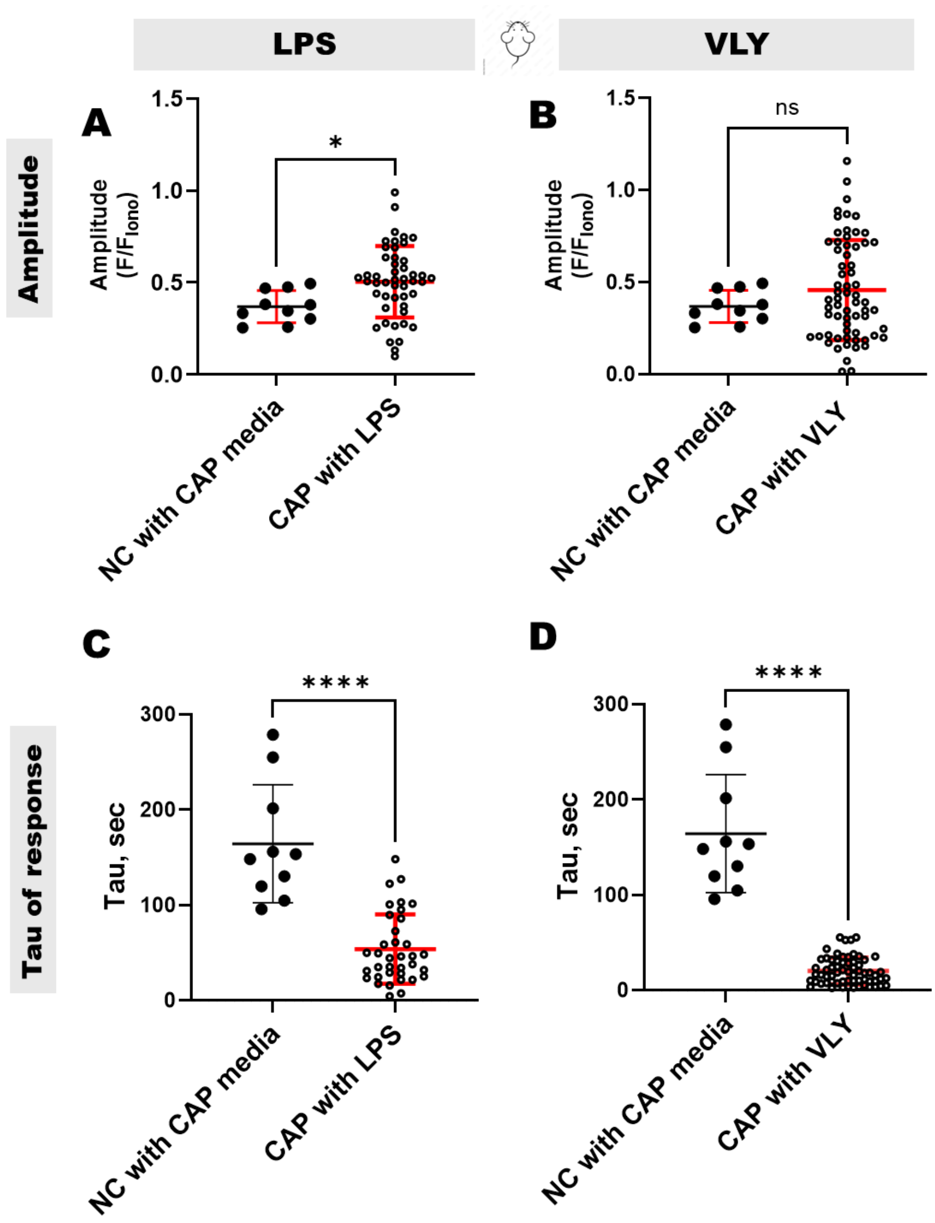


**Fig. S8. Sperm [Ca$^{2+}$]$_I$ response in WT mouse sperm is faster in the presence than in the absence of LPS or VLY. (A, B)** Amplitude and **(C, D)** tau of [Ca$^{2+}$]$_I$ response in NC sperm on acute exposure to **(A, B, C, D)** capacitating (CAP) media, or in CAP sperm on acute exposure to **(A, C)** LPS or **(B, D)** VLY. Data are presented as mean and standard deviation (n=3 biological replicates). *P<0.05, ****P<0.001 by unpaired t-test.

**Fig. S9**

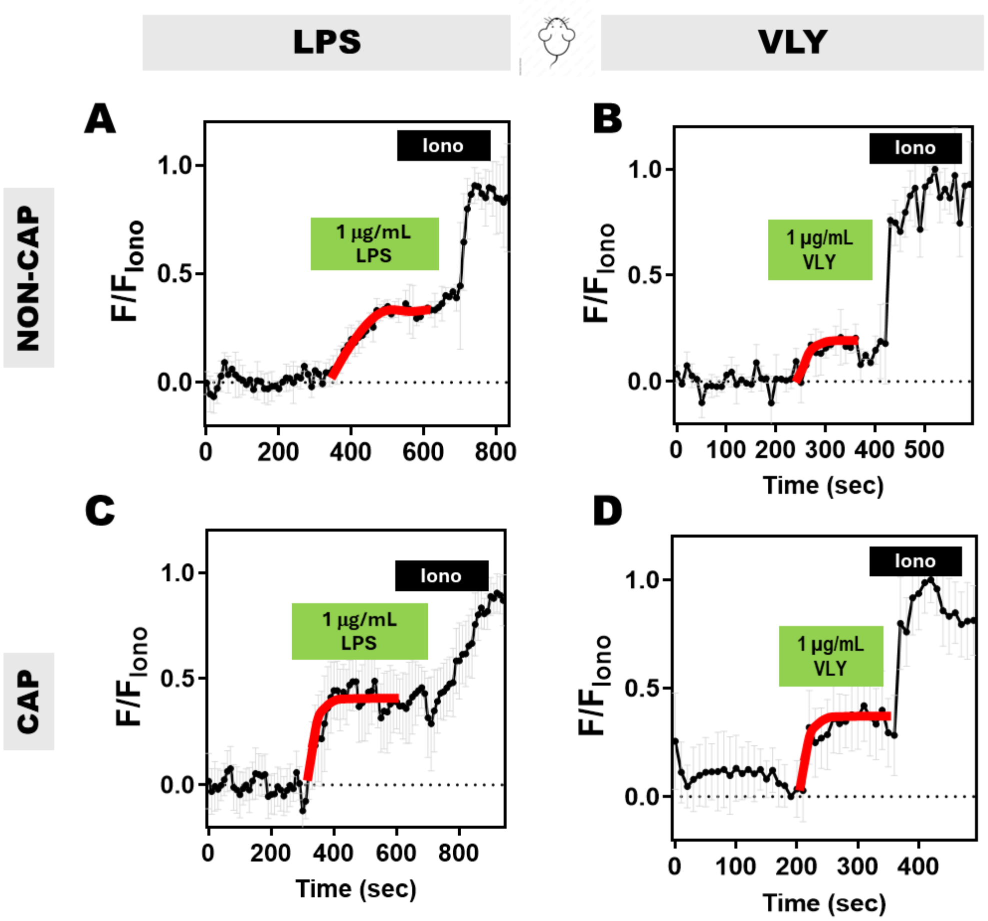


**Fig. S9. LPS and VLY induce $[Ca^{2+}]_I$ responses in mouse sperm**. Representative traces of **(A, C)** LPS- and **(B, D)** VLY-induced $[Ca^{2+}]_I$ response in (**A, B**) NC and (**C, D**) CAP mouse sperm. The red curves are standard exponential fits. Each trace was normalized to its respective ionomycin (Iono) response. Data are presented as mean and standard deviation (n=3 biological replicates).

**Fig. S10**

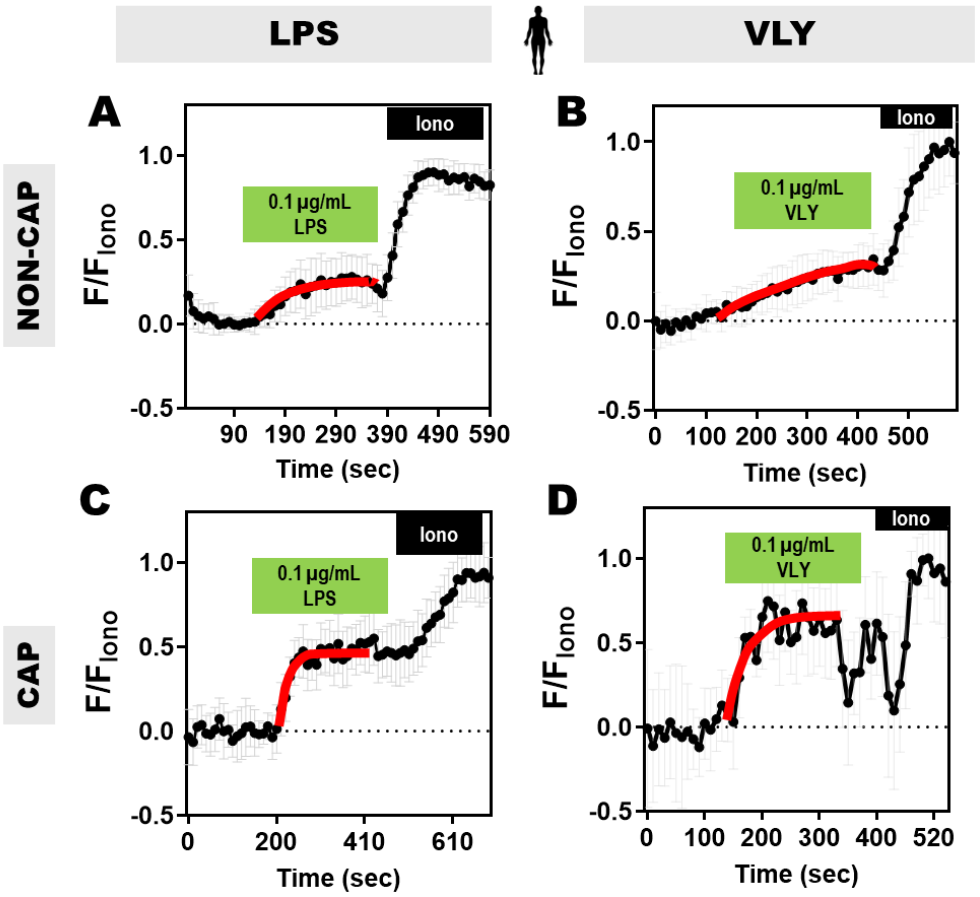


**Fig. S10. LPS and VLY induce $[Ca^{2+}]_I$ responses in human sperm**. Representative traces of **(A, C)** LPS- and **(B, D)** VLY-induced $[Ca^{2+}]_I$ response in (**A, B**) NC and (**C, D**) CAP human sperm. The red curves are standard exponential fits. Each trace was normalized to its respective ionomycin (Iono) response. Data are presented as mean and standard deviation (n=3 biological replicates).

### Fig. S11

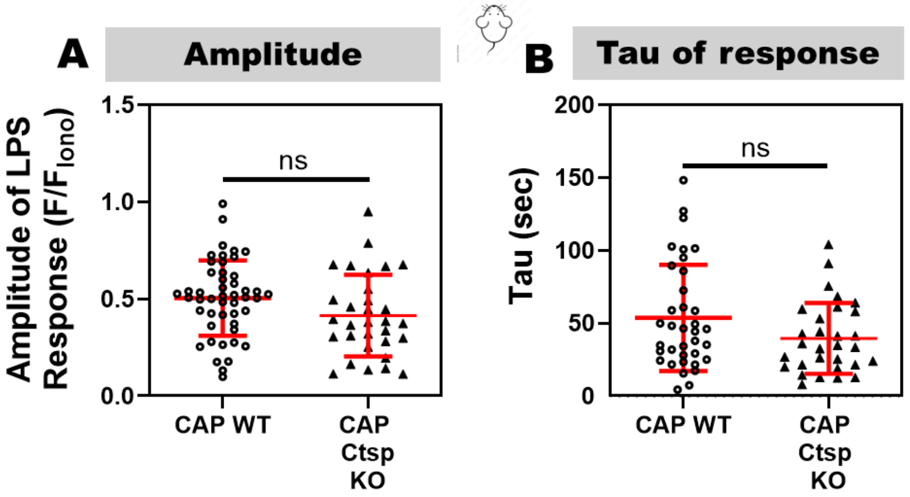


**Fig. S11. LPS-induced [$Ca^{2+}$]$_I$ increases are similar in sperm from WT and CatSper knockout mice. (A)** Amplitude and **(B)** Tau values for the 0.1 µg/mL LPS response in WT and CatSper knockout (KO) sperm in CAP conditions. Data are presented as mean and standard deviation (n=3 biological replicates). ns= non-significant by unpaired t-test.
